# New candidates for regulated gene integrity revealed through precise mapping of integrative genetic elements

**DOI:** 10.1101/2020.01.24.918748

**Authors:** Catherine M. Mageeney, Britney Y. Lau, Julian M. Wagner, Corey M. Hudson, Joseph S. Schoeniger, Raga Krishnakumar, Kelly P. Williams

## Abstract

Integrative genetic elements (IGEs) are mobile multigene DNA units that integrate into and excise from host bacterial chromosomes. Each IGE usually targets a specific site within a conserved host gene, integrating in a manner that preserves target gene function. However, a small number of bacterial genes are known to be inactivated upon IGE integration and reactivated upon excision, regulating phenotypes of virulence, mutation rate, and terminal differentiation in multicellular bacteria. The list of <u>r</u>egulated <u>g</u>ene integrity (RGI) cases has been slow-growing because IGEs have been challenging to precisely and comprehensively locate in genomes. We present software (TIGER) that maps IGEs with unprecedented precision and without *attB* site bias. TIGER uses a comparative genomic, ping-pong BLAST approach, based on the principle that the IGE integration module (i.e., its *int-attP* region) is cohesive. The resultant IGEs, along with integrase phylogenetic analysis and gene inactivation tests, revealed 19 new cases of genes whose integrity is regulated by IGEs (including *dut, eccCa1, gntT, hrpB, merA, ompN, prkA, tqsA, traG, yifB, yfaT* and *ynfE*), as well as recovering previously known cases (in *sigK, spsM, comK, mlrA*, and *hlb* genes). It also recovered known clades of site-promiscuous integrases and identified possible new ones.

## INTRODUCTION

Mobile genetic elements (MGEs) such as plasmids and prophages often carry determinants of bacterial traits such as pathogenicity, symbiosis, and antibiotic resistance (1). Transmission of these DNA units between bacterial hosts, through vehicles such as conjugation pili and bacteriophage particles, followed by stabilization in the new host, is a major mechanism of horizontal transfer and fixation of these traits. A large fraction of MGEs encode an integrase enzyme from either the tyrosine or serine recombinase protein family, that stabilizes the newly transmitted MGE by integrating it within the host chromosome. Integrases catalyze recombination between a particular DNA site (*attP*) in the circular form of the MGE and the target DNA site (*attB*) in the host chromosome, leaving the integrated MGE flanked by two *attB/P* recombinant sites, at its left (*attL*) and right (*attR*) ends (2). We introduce the term “integrative genetic element” (IGE) for this subclass of MGEs (Supp. Fig. 1) and define it as an ostensibly mobile DNA unit for which a putative *attL, attR*, and at least one contained integrase gene (*int*) can be identified. Note the stringency of this definition compared to that for the term “genomic island” (GI), which has been defined as loosely as any foreign gene cluster (3). Many IGEs bear additional gene content that subtypes them as either prophages or ICEs (integrative and conjugative elements), but for a large fraction of IGEs (measured here at ∼50%) such indicative genes cannot be identified; some of these latter IGEs may be satellites that depend on gene products from helper phages or ICEs for their presumed transmission mechanism.

Integration of an IGE into a target chromosomal gene is usually accomplished without inactivating the target gene. For example, a typical IGE targeting an *attB* within a tRNA gene carries within its *attP* a fragment of that same tRNA gene; integration does disrupt the original tRNA gene, yet simultaneously restores a functional tRNA gene, now a recombinant that incorporates the fragment from *attP* (Supp. Fig. 2). Likewise, at an *attB* target in a protein-coding sequence (CDS), the IGE typically integrates innocuously, preserving target gene function using a similar *attP* fragment strategy, or perhaps targeting an extreme tail of the protein sequence that does not affect protein function. However certain IGEs are known to control gene activity, inactivating the target CDS upon integration and reactivating it upon excision (4-15). This regulated gene integrity (RGI) can occur irreversibly in the development of nonreproducing cells of multicellular bacteria, where integrases catalyze specific IGE deletions in the chromosome that restore the *sigK* or *spsM* genes in spore mother cells (5,10,12), or the *nifD, hupL*, or *fdxN* genes in cyanobacterial heterocysts (6,8,9). RGI can also occur reversibly, as at the *Listeria comK* or the *Streptococcus mutL* genes (11,13), where excision of the IGE circle temporarily alters gene expression until the circle re-integrates (7). While many of these gene-regulatory IGEs are full-sized ICEs or prophages, others have degraded to sizes as small as a few kbp, retaining little more than the *int* gene itself (16).

Discovery of new cases of RGI must rule out the accidental gene inactivation events that can occur through occasional off-target activity of site-specific integrases, or through the action of certain clades of integrases that have become site-promiscuous. Promiscuity has emerged in multiple clades of the integrases (Y-Ints) within the tyrosine recombinase family, mobilizing IGEs such as Tn916, Tn4371, *tfs*, and CTnDOT (17-19). Promiscuity has also emerged among the serine recombinases, all members of which share a core catalytic domain; members can be further classified as either having a second larger C-terminal domain associated with IGE integrases (S-Ints) or lacking it (S-Cores) (20). This second domain has complex-stabilizing coiled-coil motifs that control S-Int recombination directionality, i.e., integration vs. excision (21). The simpler S-Core proteins were traditionally not known as integrases, but instead as resolvases and DNA invertases. However a small clade of site-specific IGE integrases has been identified among the S-Cores, the ϕRSM group (22), and a site-promiscuous S-Core clade mobilizes the insertion sequence IS607 (23).

Several bioinformatic tools have been developed to identify foreign gene clusters in genomes, based on features such as: 1) sequence composition differing from that of the surrounding chromosomal DNA, used by AlienHunter, GI-SVM, INDeGenIUS, Centroid, MTGIpick, ZislandExplorer, MJSD, SigHunt, PAI-IDA, MSGIP (24-33), 2) foreign gene content, used by IslandPath-DIMOB, SIGI-CRF, SIGI-HMM, Wn-SVM, GIHunter, PredictBias (34-38), 3) sporadic occurrence among closely related strains, used by IslandPick (39), and 4) preference to integrate into tRNA and tmRNA genes, used by Islander (40). Other packages, such as IslandViewer4 (41), amalgamate some of the above methods for improved performance.

While it is an important research goal to identify all foreign gene clusters in genomes, there is also great value in focused identification of those of the IGE class, which are more likely to be actively mobile. When the *attL* and *attR* of an IGE are properly identified, that IGE is precisely mapped. Only such precise mapping can allow proper surveys of RGI and integrase site-specificity and promiscuity. Furthermore, the association of integrases with their site specificities would enrich the stockpile of recombinases available for the biotechnology of genome editing. Features 1 and 2 above cannot inherently yield precise IGE coordinates; sequence composition rarely transitions cleanly to correctly demarcate IGE termini, while gene-based methods cannot map more finely than gene boundaries and do worse when they miss or overextend IGE terminal genes. Feature 3 allows comparative genomic approaches that could in principle map IGEs precisely, although IslandPick is gene-based with the attendant imprecision. Of all the above tools, only Islander maps IGEs precisely, by finding the tRNA gene marking one end of the IGE, and the tRNA gene fragment displaced by integration that marks the other end of the IGE; however, Islander cannot find IGEs that target non-t(m)RNA genes. Analysis of Islander IGEs presented here reveals that *int* genes tend to lie close to one end or the other of the integrated IGE (i.e., that IGE *int-attP* integration modules are cohesive (42)). This principle was used as a starting point for our new method TIGER (The IGE Retriever), which is also rooted in comparative genomics, fully exploiting feature 3 by searching for reference genomes where the integration site *attB* is uninterrupted by any IGE.

TIGER maps *attB* sites precisely and without bias, and therefore allowed us to survey site-specificity and RGI. TIGER also discovers with precision transposons and insertion sequences (ISs), which are typically mobilized by DDE transposase family members; we used this capability to mitigate a major artifact arising when TIGER is applied to search for IGEs. Additional methods recovered the previously-known promiscuous IS607, Tn916, Tn4371 and *tfs* clades, and also pointed to groups among the S-Core clade that we suspect are resolvases that are passengers on transposons mobilized by classical transposase enzymes. We have recovered known cases of RGI and discovered several new candidate cases.

## MATERIALS AND METHODS

### Datasets

Nucleotide sequence datasets used were the 2168 complete RefSeq genomes listed in Supp. Table 1 (137 archaeal, 2031 bacterial) that were used for Islander (43), 1010 temperate viruses listed in Supp. Table 2 (the bacterial or archaeal viral isolates from GenBank that we found to encode a Y-Int or S-Int), 109 experimentally-validated ICEs (44), and sets of 3266 GI-negative segments (13.8 kbp average) and 1845 GI-positive segments (11.6 kbp average) from 104 genomes, and 80 “gold standard” GI calls (38.3 kbp average) from the literature for 6 genomes (45). The software developed below, run on the 2168 genomes, yielded 6415 (TIGER/Islander mode), 4762 (TIGER-only mode) and 3191 (Islander-only mode) IGEs. Mock IGEs (32075) were prepared by taking a segment from a random location of the same source replicon and of the same length as each TIGER/Islander IGE, five times.

### Genome processing

The TIGER software package is available at github.com/sandialabs/TIGER. Its script tater.pl handles annotation of raw multi-FASTA genome sequence files. Prokka 1.11 (46) is called as “prokka --rfam --prefix protein --gcode Genetic_Code --kingdom Kingdom -- cpus 1 --rnammer --notrna --outdir ./ --force --quiet --locustag Nickname genome.fasta”, where the genome-identifying Nickname is a three-letter genus/species abbreviation followed by a serial number, and Kingdom (Archaea/Bacteria) and Genetic_Code are taken from the assembly report file and taxdump.tar file from NCBI. tRNA/tmRNA genes are carefully reannotated with the tFind.pl module (43) (manuscript in preparation). Islander was run as described (43). Pfam-A (47) and other HMMs noted below were applied to all protein sequences using hmmsearch (HMMER 3.1b2, http://hmmer.org/), collecting the top above-threshold match. Transposase genes were called using a set of Pfam and other HMMs as described (48). Three Pfam HMMs were used to call integrase genes: Phage_integrase (Y-Ints), Resolvase (the catalytic domain shared by S-Ints and S-Cores) and Recombinase (the second domain unique to S-Ints). Certain Phage_integrase members serve roles other than IGE integrases, such as the Xer housekeeping proteins that resolve chromosome dimers after replication and the IntI integrase that mobilizes integron cassettes. From among the Phage_integrase matches, Xer proteins were identified using four Xer subfamily profiles from HAMAP as described (43), and IntI proteins were identified using the intI_Cterm HMM as described (49).

### TIGER core module

The tigercore.pl module aims to identify and map IGEs by finding reference genomes in which the IGE integration site is uninterrupted (i.e., contains no IGE). It takes three main inputs: a scaffold/replicon DNA sequence, a coordinate on that DNA (here, the midpoint of an integrase gene), and a reference genome BLAST database (here, refseq_genomic downloaded from NCBI on April 4, 2017). Two query sequences (q1L and q1R) of 15 kbp are taken from the replicon, to the left and right of the coordinate, and used to probe the database with BLASTN in default mode [Fig. 2]. Matches longer than 500 bp are processed further, filtering out those that fully reach the input coordinate (indicating reference genomes that contain the same putative IGE). For each match, a return query of 3 kbp is taken from the reference genome region adjacent to the coordinate-proximal end of the match, reaching back into the matching region 250 bp to include the direct repeat (DR) sequence. The set of return queries (q2) are used with BLASTN against the replicon to find the matching distal flank of the IGE.

**Fig. 1.**
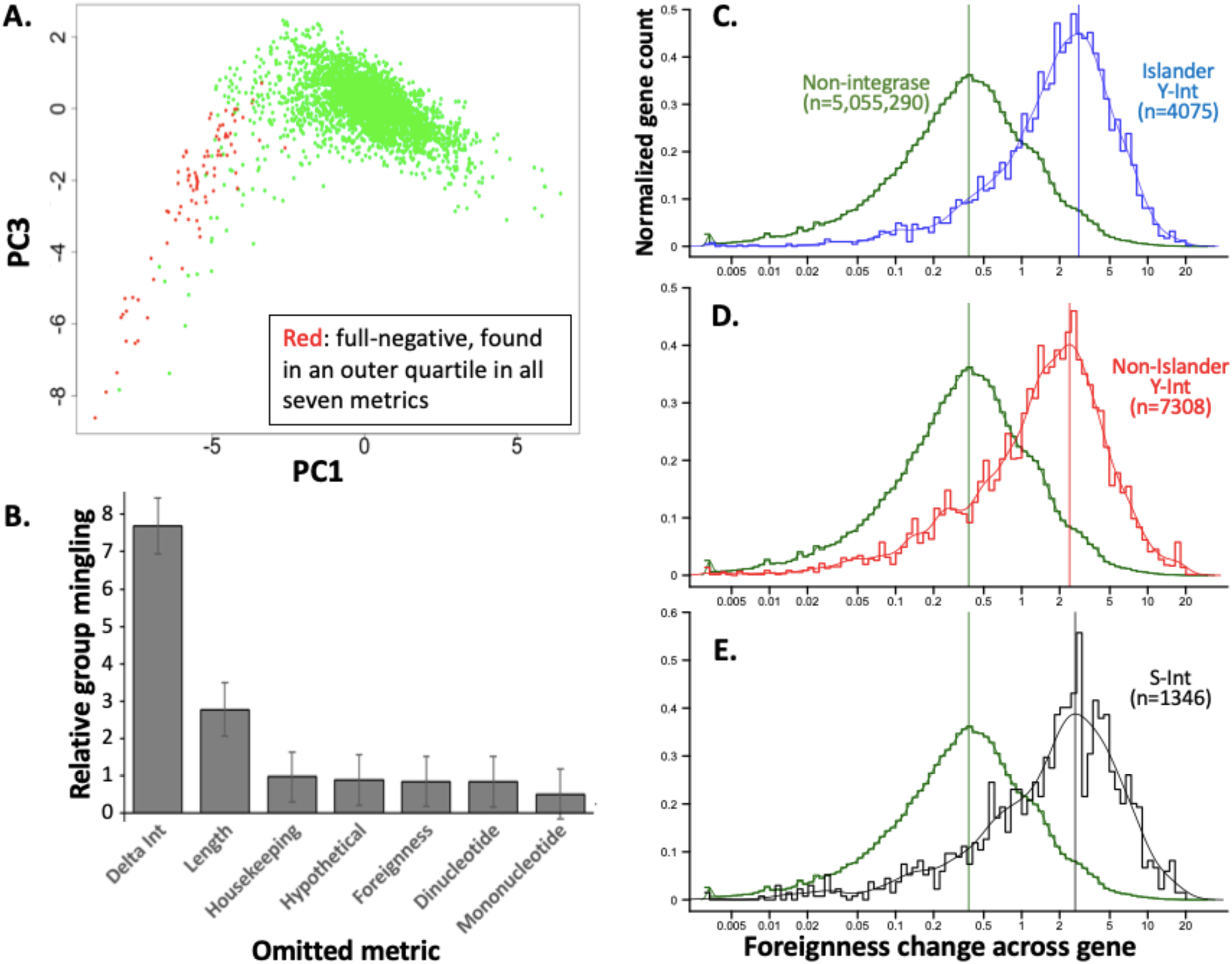
Promise of the integrase module cohesion (delta-int) principle for IGE discovery. **A**. Horn of full-negative IGEs. Islander IGEs found in an outer quartile for all seven IGE metrics (Supp. Fig. 3) are marked in red in principal components analysis of those seven metrics. PC1 vs PC2 was similar, but PC1 vs PC3 spread the data more and was therefore selected for visualization. **B**. Robustness of each metric. Multidimensional distance was taken for each cross-group point pair, from the full-negative group and the remaining group, for all seven metrics (D_original_) or after omitting each metric in turn (D_leaveout_). Mingling of the two groups was calculated as the average of 1-(D_leaveout_/avg. D_original_) values for the cross-group point pairs; these values were normalized to total data spread (mean of total pairwise distances of all point pairs). Error bars mark standard deviation. **C**. Foreignness change at Islander Y-Int genes. Foreignness was pre-evaluated for each Pfam category using a set of reference plasmid and phage genes. For every protein gene among 2168 study genomes, windows of three neighboring Pfam genes were taken from its left and right side, and foreignness change was taken as absolute difference in summed foreignness for the two windows (note logarithmic scale). **D**. Promise for IGE discovery at non-Islander Y-Int genes. **E**. Promise for IGE discovery at S-Int genes.

**Fig. 2.**
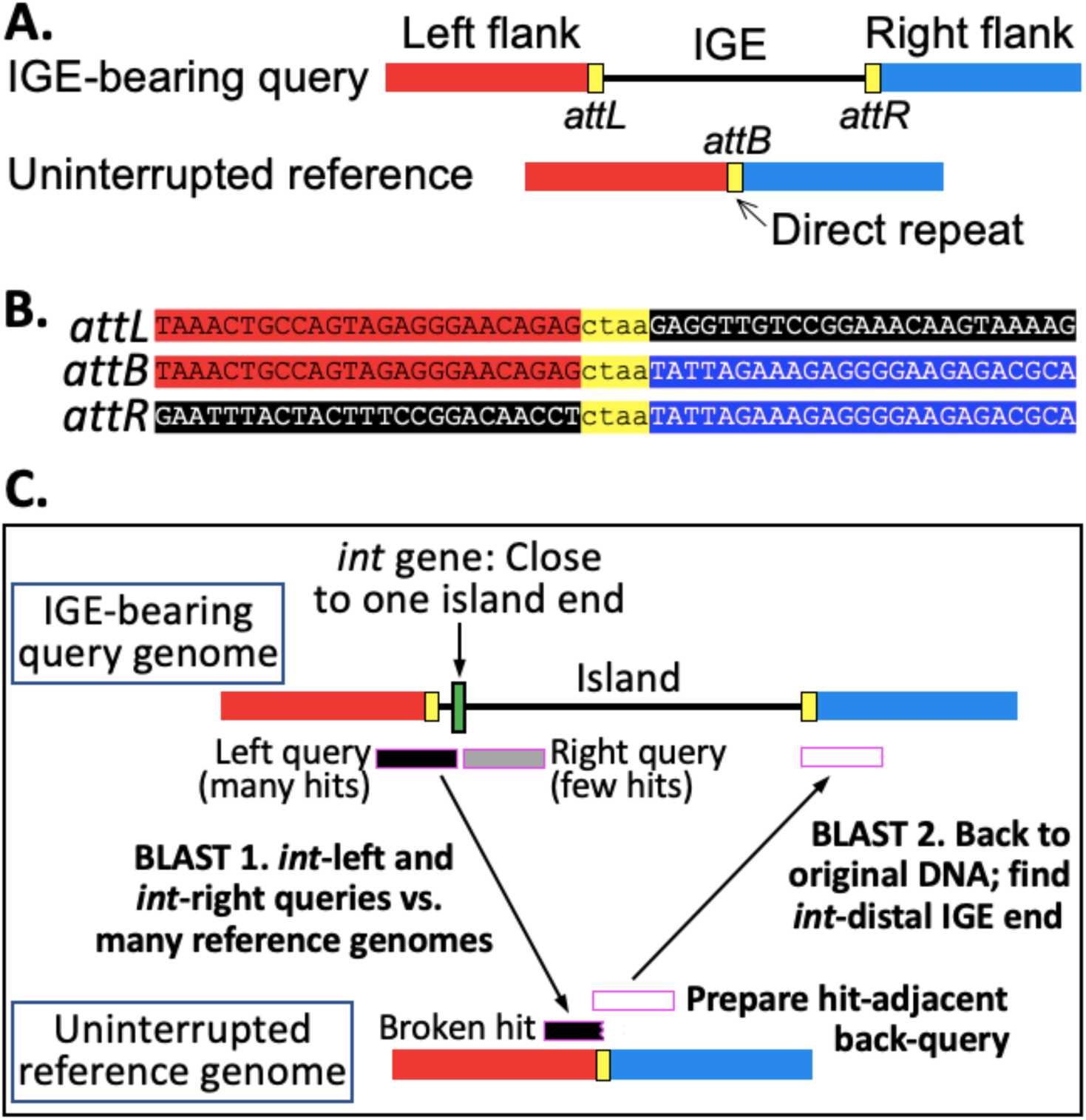
TIGER: ping-pong BLAST for IGE discovery. The corresponding regions of an IGE-bearing and uninterrupted reference genome pair (**A**) produce a sequence alignment pattern (**B**). Strand crossover presumably occurs with the direct repeat block (yellow). In TIGER (**C**), the first BLAST simultaneously locates the *int*-proximal end of the IGE and the *attB*, and the second locates the distal end of the IGE.

### TIGER finishing module

Successful matches from above are tested for the IS artifact, i.e., whether a q1 BLAST match lies entirely within a transposon. If a transposase gene is located in the q1 match, the boundaries of the potential transposon are probed by running the core module in IS mode using the midpoint of the transposase gene as the coordinate, a DR reachback of 30 bp, and IS size limits of 5 and 15 kbp. If no IS is determined by TIGER, a second search employs BLASTN of the q1 match against the ISFinder database (50). If either approach shows that a q1 match lies entirely within an IS call, the IGE call is rejected. This module also collects support values for each IGE call, i.e. the number of uninterrupted reference genomes identified. Sequence variations in the *attB* vicinity among reference genomes may cause slight fluctuations in the termini of BLASTN matches. If so, the resultant set of slightly variant IGE calls are merged, applying the total of their support values to the best-supported call among them. Duplicate calls from both q1L and q1R for short IGEs are deduplicated. Gene target information is taken from the annotation of a supporting reference genome if available, otherwise from the annotation of the query genome.

### False positive metrics

Calculation methods are detailed here for seven metrics whose utility in identifying false positives is demonstrated in Supp. Fig. 3 and described in Results: 1) Mononucleotide bias is the absolute difference between the relative abundances (51) of G+C in the candidate IGE and its scaffold/replicon. 2) Dinucleotide bias is taken as described (51). A Pfam enrichment factor table was prepared for the 3) housekeeping metric, noting the top Pfam match for each gene among the study genomes; for each Pfam, the enrichment factor was taken as its count among the Islander IGE set genes divided by its count among all genes. Housekeeping for a candidate IGE is taken as the average of enrichment factors for its genes with Pfam calls, subtracting the corresponding value for all genes in its scaffold/replicon. 4) Foreignness is taken as for housekeeping, except that the Pfam enrichment table was prepared using a set of plasmid and phage genomes (Supp. Table 3) instead of the Islander IGEs. 5) Hypothetical is the fraction of IGE protein genes with no Pfam calls, subtracting the corresponding value for all protein genes of the scaffold/replicon. 6) Length is for the IGE in bp. 7) Delta-int is the shortest distance in bp between an *int* gene in the IGE and a terminus of the IGE, replaced with one if an *int* gene extends beyond an IGE terminus.

### Islander false positive score formula

Onto a display of principal components for Islander IGEs based on the seven false positive metrics described above (Supp. Fig. 4B), a convex hull was drawn around IGEs confirmed by TIGER. This defined IGEs for training purposes as either positive (within-hull) or negative (outside-hull). Using a randomly chosen 75% of the data, a generalized linear model was trained to predict positives and negatives using the seven metrics. Specifically, the caret module in R was used with default options to train a model using the “glm” method, which uses the basic R function glm. Model performance was tested using the remaining 25% of the data, yielding an area of 0.97 under the receiver operating characteristic curve (where 1 would indicate perfect performance). Coefficients were extracted from the generalized linear model, generating a formula for identifying false positives (Supp. Fig. 4D).

### Resolution module

This module handles relationships between IGE calls within a genome, and operates in Islander-only, TIGER-only or combined (described here) mode. The termini of raw Islander calls are collected. This list of termini is grown by adding only those raw TIGER calls that share a terminus with an Islander call. Termini are grouped as much as possible into tandems, by the IGE calls that connect them pairwise. These Islander/TIGER mixed tRNA gene tandems are resolved at the internal termini into IGE units, insisting that each IGE have at least one *int* and respecting lower and upper size limits (2 kbp and 200 kbp). IGEs are not allowed to overlap; such conflicts are resolved by rejecting the call with the lower TIGER support value, otherwise with the higher false positive formula score. Then TIGER-only tandems are built similarly from the remaining raw calls and resolved into IGE units, and all tandems are de-overlapped. Either tandem resolution step can produce IGE calls that have no direct TIGER support but are instead inferred. All calls failing the false positive formula cutoff are rejected, as are all integrases failing the cutoff formula for top:second-best support ratios (Supp. Fig. 4) and the IGE calls that depend on them.

### Deduplicating IGEs

Because some genomes in this study are very closely related, there may be nearly identical IGEs among our set due to vertical inheritance after a single integration event in an ancestor. Such vertical inheritance would confound our analysis of promiscuous and Pfam-disrupting integrase clades. To address this issue, we grouped IGEs into similarity clusters and deduplicated. Pairwise average nucleotide identity (ANI) values between IGEs were taken using fastANI (52) with arguments “-k -t 120 --fragLen 500 --minFrag 2” to accommodate the minimum size allowed for our IGEs. The resulting distance matrix was converted into a network and filtered for IGE pairs that had ANI > 0.95 and alignment fraction > 0.9. The Floyd–Warshall algorithm, which identifies the shortest path between nodes in a weighted graph and was implemented from the Python SciPy library, was applied to the remaining IGE pairs to calculate the lengths of shortest paths between all pairs of IGE. The result was clustered using DBSCAN (from the python module sklearn), with eps (radius) = 1 and min_samples = 2. Cluster members were replaced with a single representative for each cluster.

### Integrase phylogenetic analysis

Sequences of integrases that were unique in an IGE were aligned using hmmsearch (HMMER 3.1b2, http://hmmer.org/) with the -A flag against the Pfam HMMs PhageIntegrase (Y-Ints), Recombinase, and Resolvase, concatenating the Resolvase and Recombinase alignments (S-Int/S-Core) to reunite the two separately processed domains of each S-Int protein. Trees were prepared for the two families using double-precision FastTree 2.1.10 with the LG substitution matrix (53), which for both families produced higher likelihood scores than the WAG or JTT matrices. Additional trees were built in parallel that added numerous reference family members (145 Y-Ints and 86 S-Int/S-Core, including 46 IS607 integrases from ISFinder) allowing annotation of clades in the IGEs-only trees. An HMM was built from a Muscle alignment of the reference IS607 integrases that, when applied to our IGE and reference integrases with a cutoff bit score of 64, perfectly collected all and only the IS607 clade members identified phylogenetically. Bootstrap sets (1000) of each of the two original alignments were taken using Phylip, and trees built as above.

### IGE subtyping

The script polish.pl types IGEs. Phage calling was based on lists that we curated of Pfam HMMs enriched in bacteriophage proteins, categorized as either virion or non-virion, omitting those producing significant cross-talk with ICEs or GI-negatives (Supp. Table 4). Two heuristics were devised for phage calling: Phage1 (query matches at least one virion and one non-virion phage HMM), or Phage2 (query matches ≥1 phage HMM from one of these categories). ICE calling employed a set of 47 HMMs for proteins functioning in ICE replication or DNA transfer, omitting those for integrases and FtsK_SpoIIIE (44). Two heuristics were devised: ICE1 (query has ≥7 ICE HMM matches that account for at least 15% of its proteins), or ICE2 (query has ≥2 ICE HMM matches, accounting for ≥12% of its proteins if <10 kbp or ≥7% of its proteins if ≥10 kbp).

Resolution rules for ICE/phage double calls were based on inspection of the 26 instances found among TIGER/Islander, temperate phage, and ICE datasets. Phage1/ICE double calls were found to be either an unresolved ICE/phage tandem or an IGE of one type integrated within another (54), yielding a new category of IGE call (ICE+Phage). Phage2/ICE double calls were converted to the ICE call. Supp. Fig. 10 shows performance of this system on temperate phage, ICE and IGE-negative references. Results for the temperate phage, ICE, and IGE-negative datasets (with ICEs serving as negatives for phages, and vice versa) allowed calculation of recall and precision for phage calling as 98.2% and 99.7%, and for ICE calling as 75.2% and 98.8%.

## RESULTS

### Islander false positives highlight the cohesion of the integration module in IGEs

We improved the initial genome preparation steps of Islander (43), particularly the annotation of DNA recombination genes: S-Ints, S-Cores, Y-Ints, integron integrases, Xer resolvases, and transposases. Islander was applied to a study set of 2168 complete bacterial and archaeal genomes. Islander employs several filters to remove false positives, but these are not completely effective. We devised seven metrics for evaluating false positives, six of which have been used by others to distinguish IGEs from core chromosomal segments, based on: nucleotide composition (Mononucleotide bias, Dinucleotide bias), gene content (Housekeeping, Foreignness, Hypothetical) or other criteria (Length, Delta-int). Delta-int is defined as the shortest distance between an integrase gene in the IGE and a terminus of the IGE, employed because we and others have noted that *int* genes typically lie near an end of the IGE. Each of the 3191 Islander IGE calls was scored by these metrics, and each metric tended to form a distribution with a single peak, except Length which had two peaks, at ∼10 kbp and ∼45 kpb (Supp. Fig. 3). We found 79 IGE calls that were in an outer quartile for all seven metrics, identifying a set of full-negative calls. Furthermore, these full-negatives were all in the same outer quartile in each metric, identifying which side of each distribution might best identify additional false positives.

The full-negatives clustered in one horn of a plot of principal components (Fig. 1A). This clustering was measured by taking the average 7-dimensional Euclidian distance between all cross-group point pairs, for the full-negative group and the group of remaining points. Seven leave-out experiments were performed, omitting each metric and measuring how intermingled the full-negatives became among the other IGEs, as the average pairwise 6-dimensional distance (Fig. 1B). The most disruptive leave-out (i.e., the most important for distinguishing full-negative from other IGEs) was delta-int. Delta-int is effectively a measure of cohesion of the integration module (42), and its importance reflects the evolutionary principle that DNA-active enzymes should be located near the DNA site (here, *attP*) that they act upon. Gene content and nucleotide composition leave-outs had smaller effects.

### Promise of other integrase genes for marking IGE termini

Having demonstrated the importance of integration module cohesion in Islander IGEs, we considered that it might serve as a broad IGE-finding principle, reasoning that an *int* gene tends to mark a terminus where IGE sequence transitions to chromosomal sequence. We measured the gene foreignness change (Methods) across Islander integrase genes as 8.1-fold higher than for non-integrase genes (Figure 1C).

Islander insists that its output IGEs contain a Y-Int, a class known to favor integration sites within tRNA genes. Non-Islander Y-Int genes likewise had a much (6.9-fold) higher foreignness change than non-integrase genes (Figure 1D), showing that the principle has promise for finding additional IGEs, which may reside in sites that are not in tRNA genes.

A smaller number of IGEs use a member of the S-Int group, which has not been known for targeting tRNA genes. To assess whether the integration module cohesion principle also has promise for finding IGEs that use this other main integrase group, foreignness difference was measured across S-Int genes. Indeed, these genes also have a much (7.6-fold) higher foreignness difference than non-integrase genes (Figure 1E). These results indicate that *int* genes generally mark the boundary between chromosomal and foreign DNA segments.

### TIGER algorithm

The integration module cohesion principle described above can approximately locate one end of an IGE. To better locate this *int*-proximal end, as well as the *int*-distal end, we added a second principle for detecting IGEs, their sporadic occurrence among closely related genomes. By identifying reference genomes where the putative IGE integration site is uninterrupted, sequence matching tools can precisely locate both termini of the IGE. The number of uninterrupted reference genomes found for an IGE is a measure of its comparative genomic support. TIGER applies two phases of BLASTN, first using queries from the left and right of the *int* gene midpoint against a reference genome database, then querying back from each matching reference genome to the original *int*-bearing genome (Fig. 2). The desired phase I match should not be full-length (which might indicate a genome containing the same IGE), but should be truncated at the *int* end of the query, terminating at the actual junction between chromosomal and foreign DNA. The location of this truncated match determines the phase II query, which overlaps the match somewhat to catch the typical direct-repeat block (yellow in Fig. 2) found at the two ends of the IGE. For tRNA genes this block is typically <100 bp, but it became necessary to raise the overlap length to 250 bp to accommodate some of the longer direct repeat blocks found for some protein-coding gene targets, like *icd*. Minimum (2 kbp) and maximum (200 kbp) IGE size limits were imposed. Matches to a candidate IGE may arise from multiple uninterrupted reference genomes; the number of such genomes is taken as the TIGER support value for the IGE.

IGE nomenclature follows the formula Genome.Size.Target, with the genome designated by a three-letter genus/species abbreviation and a serial number, size in kbp and rounded off; see Supp. Table 8 legend for *attB* naming conventions.

### Utility in detecting Iss

Implementing this algorithm, the most notable artifact arose from IS mobility. In particular, phase I may match only an *int*-flanking IS that occurs at a different nearby site in the reference genome. This artifact can manifest as a series of IGE calls for the same integrase, with one terminus fixed and the other (corresponding to the IS-containing end) varying over a large range for different reference genomes with the IS transposed to different loci. To help resolve this artifact we discovered that TIGER can also effectively map ISs, using a transposase gene as the seed rather than an *int* gene, and adjusting minimum, maximum and overlap lengths. The ability to detect IS matches was augmented using data from the ISfinder website (50). The artifact was effectively controlled using these methods to identify and reject IGE calls if either the phase I or II matches is limited to an IS. Global IS discoveries by TIGER will be presented in a later publication, but we validate its ability to discover ISs in a study of the *E. coli* MG1655 genome (Supp. Table 5).

### Formula for false positives

TIGER analysis confirmed a large number (∼40%) of Islander calls, from the main cluster and apart from the full-negatives (Supp. Fig. 4A). A convex hull was traced around these doubly-confirmed calls (Supp. Fig. 4B), separating Islander calls into within-hull and outside-hull categories (Supp. Fig. 4C). Linear regression with the seven metric scores produced a formula (Supp. Fig. 4D, Methods) and cutoff value that optimized separation of the two categories (Supp. Fig. 4E) and that we used to automate false-positive detection. The formula left no calls with overlap to IGE-negative genomic regions (45) (Supp. Table 6).

Although many false positives were rejected this way, some interesting cases were also lost. For example the raw call Xor4.60R (NC_017267.1:4654483-4594551) had high TIGER support (220) and was confirmed by Islander but was rejected by the formula due mainly to its gene content that included the only genomic copies of essential *glySQ* genes. Inspection suggested that an original IGE at a tRNA-Arg gene had been invaded by a transposon formed from two IS-Xo4 copies flanking the *glySQ* region.

### Recombination activity not due to the candidate integrase

Other mechanisms besides the action of our candidate integrases may result in recombination events that lead to false positive IGE calls. Negative or other selective pressure on genes in the vicinity of an *int* may drive multiple such non-integrase recombination events among reference genomes, each perhaps occurring at different termini. We observed cases of multiple overlapping calls where the second-best support value was not far from the top support value. One case that was egregious and simple to understand was the signal around the integrase for the well-known IGE PLE5, one of a group of *Vibrio* satellite prophage “PLEs” (<u>p</u>hage-inducible island-like <u>e</u>lements), all of which are induced by the lytic phage ICP1 (55). PLE5 is found integrated into the superintegron of *Vibrio cholerae* O395, using one of the superintegron cassette attachment sites (*attC*) as its *attB*. The PLE5 IGE was nested among 58 different raw TIGER calls; the top call had a support value of 15 and the second-best support was five, only 3-fold lower, while the actual PLE5 was called but with a support value of four (Supp. Fig. 5). The numerous calls overlapping PLE5 have termini in the many superintegron *attC*s, indicating that they are due to the distally-encoded integrase IntI of the superintegron, not the PLE5 integrase. In other cases with low top:second-best support ratios, the presumed alternative recombination system may be less readily identified, perhaps falling under the umbrella of illegitimate recombination.

We used the top:second-best support ratio and the number of overlapping raw TIGER calls to reject *int*s that may be subject to such non-integrase recombination events, and thereby reject any IGE calls that depend on them. Supp. Fig. 6 shows a scatter plot of these data for the 2510 *int*s that had more than one overlapping IGE call (5148 *int*s had a single TIGER call). From discontinuities in rejected IGE statistics (Supp. Fig. 7), a cutoff was chosen that rejected 4.6% of the IGEs that would otherwise have been retained, but only 2.1% of the doubly-confirmed IGEs.

Additional *Vibrio* superintegrons had IGEs, again with high signals suspected to arise not from the IGE integrase itself, but from the normal action of IntI on integron cassettes. For each of these we manually identified the likely primary IGE between adjacent *attC*s: one was a PLE4 relative and the other four were IGEs unrelated to any PLEs (Supp. Table 7).

### Tandem IGE arrays and overlap resolution

Another challenge for TIGER and IGE-finding in general is tandem arrays (56). Typically, integration of an IGE into an uninterrupted site restores the core of integration site (the strand crossover region) at both ends of the IGE. A second IGE may then integrate into either of these, and so on, producing tandem arrays of diverse adjacent IGEs. These tandem arrays may be important for IGE evolution as within-array recombination events between chromosomal neighbors may create hybrids, as appears to be responsible for the monophasic nature that partly defines *Salmonella enterica* serovar Typhi (56). When TIGER calls overlap, we use the support values (number of uninterrupted reference genomes) to resolve them. One challenge to resolution is depressed support values for individual IGEs within arrays, as fewer (perhaps even zero) reference genomes may have the precise combination of left and right flanks for each IGE because increasing numbers of IGEs in an array tends to depress support values for each; fewer (and sometimes zero) reference genomes may have the precise combination of the IGE’s left and right flanks. Further bias against finding reference genomes that map particular IGEs in an array can come from differing preferences of each IGE’s integrase for the recombinant *att* sites along the array, as has been shown for IGEs that target the *icd* gene (57).

We prepared resolution software that seeks to build and then resolve tandem arrays. It further resolve cases of overlapping raw IGE calls, applying the false positive formula and negative selection cutoffs. It runs in Islander-only, TIGER-only, or Combined modes. The Combined mode yielded 6415 IGEs (Supp. File 1). Many were supported by both Islander and TIGER, but each method produced an even larger number of method-unique IGEs (Fig. 3A). Thus, combining the two approaches yields a more comprehensive IGE set.

**Fig. 3.**
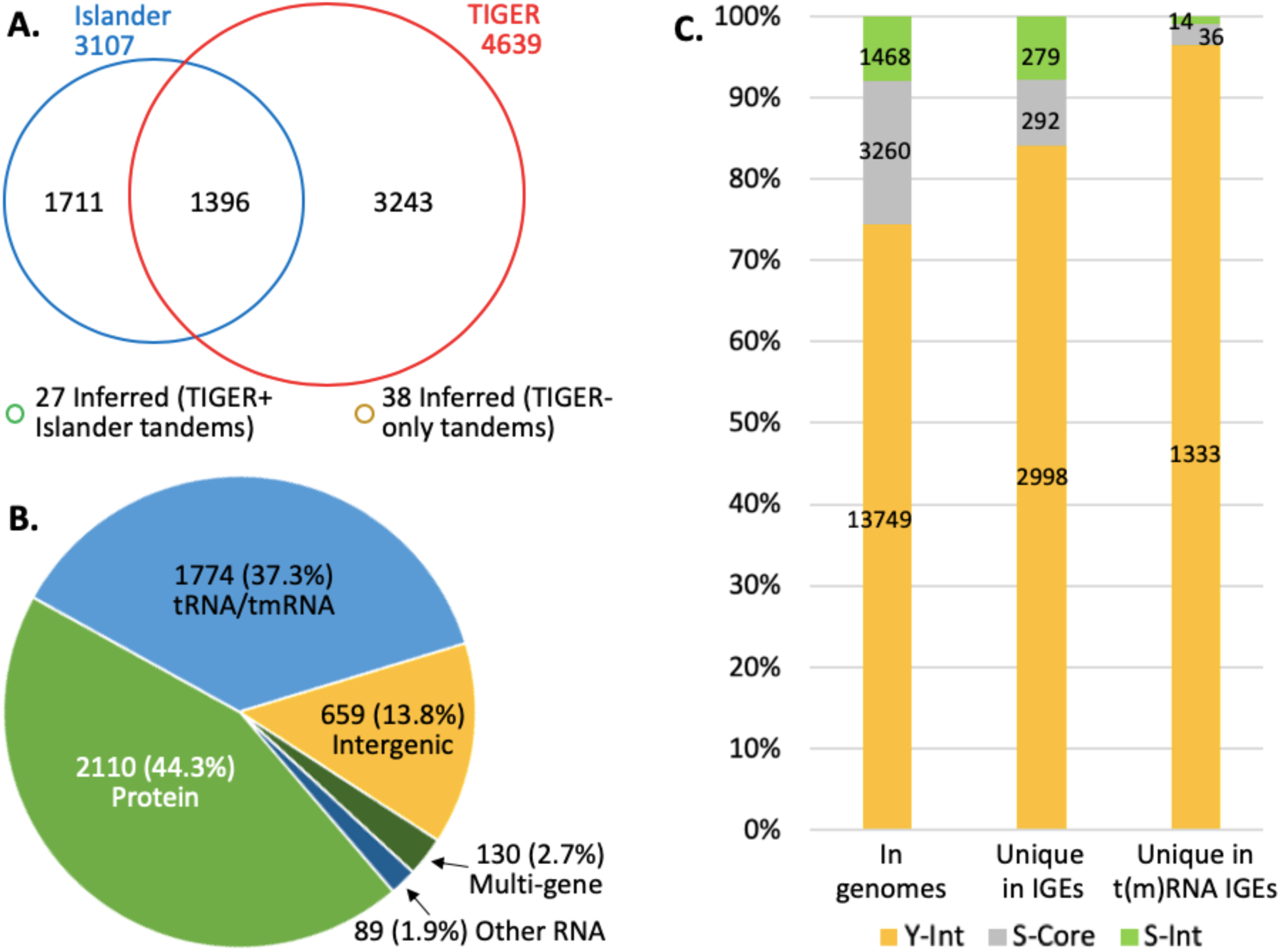
TIGER output. **A**. IGE yields for 2168 genomes from TIGER/Islander Combined-mode resolution. **B**. *attB* site types from TIGER-only-mode resolution. “Multi-gene” indicates that the direct repeat segment at *attB* overlaps two or more genes. For comparison, our annotations showed that protein genes occupied 87.1% of the DNA in these genomes, t(m)RNA genes only 0.15%, other RNA genes 0.67%, and intergenic spaces 12.0%. **C**. Integrase usage by IGEs with unique *int* candidates from TIGER-only-mode resolution.

A taxonomic breakdown of IGE counts (Supp. Fig. 8) shows Gammaproteobacteria and Acidithiobacillia at the top, averaging nearly 6 IGEs per genome. Raw support values vary taxonomically, roughly trending with the number of phylogenetically close reference genomes available (Supp. Fig. 9); to counter this effect we also present support percentages normalized to the top support value for the genome.

### Surveying attB locus type and integrase family usage

TIGER, unlike Islander, has no inherent bias for integrase family or for target site, and thus surveys these attributes. Resolution software run in TIGER-only mode yielded 4762 IGEs; the genomic feature types of their *attB*s are shown in Fig. 3B. The preference of integrases for t(m)RNA genes has been noted previously (58,59); here we measure 37.3% of IGE *attB*s in t(m)RNA genes, 248-fold higher than the 0.15% of the total genome space occupied by such genes. Among the most frequently-used protein-coding gene targets, several have been described before in the literature, while many are new discoveries (Supp. Table 8). New RNA gene targets are also indicated; e.g., we find 13 *ssrS* genes encoding the 6S RNA interrupted by IGEs (that restore the gene), in both gram-negative and -positive species.

Several TIGER-only IGEs (1176, or 24.7%) had multiple candidate *int* genes. Excluding these ambiguous cases, IGEs use *int* types at approximately the frequencies that they occur in genomes (Fig. 3C), with the notable exception that a lower fraction of S-Cores are used. The dominance by Y-Ints is extreme at t(m)RNA gene sites. For those TIGER IGEs with a unique integrase, the t(m)RNA gene usage rate is 1333/2998=44.5% for Y-Ints, significantly higher than 14/297 = 4.9% for S-Ints, *χ*^2^ (1, *N* = 3295) = 176.7, *p* = 3e-40. Use of a t(m)RNA gene by an S-Int has not been reported previously to our knowledge, but because the number is so low, experimental proof will be required.

### IGE type

The two main recognized types of IGEs are prophages and ICEs. We developed a custom system to call these types for bacterial IGEs based on content of phage/ICE protein genes (Supp. Fig. 10). Control sequence sets were for temperate phage isolates, ICEs, mock IGEs, and IGE-negative genomic segments. This system yields stronger (Phage1) and weaker (Phage2) phage calls, the latter including filamentous prophages and perhaps trailing off into defective prophages. The Phage1:Phage2 ratio was much higher (10.9) for the phage isolate controls, which presumably had been validated by plaque formation, than for IGEs (2.71), suggesting that weak calls tend to indicate unproductive degraded prophages, or satellites that cannot produce plaques on helper-free hosts. Even the strong calls contain some defective phages; of the eight *int*-containing regions annotated as “cryptic prophages” in the *E. coli* MG1655 genome (all confirmed by TIGER), four were called Phage1 (one other was called Phage2, and three did not score as phages or ICEs). Correcting for the recall rates with the positive controls, we estimated that 38.5% of IGEs are prophages and 12.2% are ICEs, leaving 49.3% with an uncharacterized mobility mechanism, perhaps largely satellites or degraded forms of the main types.

### Benchmarking

Benchmarking is challenging because of the lack of broadly accepted gold standards, the diversity of principles upon which software is based, and differing definitions of study objects. Bertelli et al (45) have prepared sets of GI-positive and GI-negative segments from 104 test genomes, and 80 gold standard genomic island calls from 6 genomes, with scripts for comparing software performance. We note that their definition of GIs (any foreign gene cluster) is substantially broader than that for IGE (Supp. Fig. 1), such that we expect TIGER to miss some of these GIs. We applied this analysis system to TIGER (Supp. Table 9), and it had the highest precision of the 21 tested programs, for both the larger and gold standard datasets, followed in both cases by Islander. “Precision” is meant here in the information retrieval sense, and as implemented indicates that TIGER results included none of the GI-negative regions. As expected due to the broad GI definition of the test system, recall was more moderate for TIGER, ranking eleventh for the larger dataset and seventh for the gold standards.

### Revisiting gold standards

Bertelli et al (45) use six “gold standard” genomes for their GI evaluation system; we propose adding a seventh genome, that of *Lactococcus lactis* IL1403, because of the exceptional work of Chopin et al (60), who identified *attL*s and *attR*s for six prophages, and successfully verified five of these by inducing, plaque-purifying, and sequencing PCR products from the DNA circularization junctions. We compare the resultant set of 86 literature GI calls to the 63 IGE calls of TIGER for these same seven genomes, to generate a new gold standard list more stringently defined as IGEs (Supp. Table 10).

We grouped the well-matched TIGER and literature GI gold standards into a new gold category of 50 IGEs, correcting the genome coordinate anomalies for those that had been determined by gene-based methods (discrepancies in other cases are due to different ways to report the direct repeats in the *att* sites). We report a second “silver” tier of suspected IGEs, with stronger disagreement between literature and TIGER calls; these all have convincing *att* sites (Supp. Table 11) and other reasons to favor the available TIGER calls, although users may wish to await experimental proof. Finally, 33 calls were dismissed as suspected or outright non-IGEs in cases that either contained no *int* gene, or in which the authors searched for but did not find *att* site direct repeats; some may have been IGEs that suffered chromosomal rearrangements that remove one *att* site region.

Six TIGER failures were revealed in this analysis: three very low-support false positives and three false negatives. Two of the latter were present among raw TIGER calls but improperly omitted from the final TIGER list. One such omission was the well-studied SopEphi prophage, which is usually found integrated into the tmRNA gene in *Salmonella* strains, but in *S. enterica* Typhi is found at a distant site within another IGE (Sen346.120.F, itself integrated into a tRNA-Phe gene). SopEphi in *S. enterica* Typhi nonetheless bears in *attL* a fragment of the tmRNA gene (56), suggesting that its location may represent an off-target insertion event. The resolve.pl module of TIGER disallows such a nested, IGE within IGE, configuration.

### Promiscuity

The above work yielded thousands of cases where an integrase protein sequence was mapped to the DNA site it presumably had used as an *attB*. Building a phylogenetic tree for those integrases then allows a search for clades exhibiting DNA site-promiscuity. We hypothesize that a promiscuous clade should exhibit diversity of *attB* usage yet be relatively shallow (with little-diverged integrase protein sequences). To reduce bias from vertical inheritance, 1622 of the TIGER/Islander IGEs were placed into 505 similarity clusters and deduplicated, replacing them with a single representative of each cluster (Methods). Additional *int*-ambiguous IGEs were excluded from analysis if among the 1521 with multiple nominal *int* genes. This left 3978 Y-Int and 465 S-Int/Core sequences whose principal domain sequences were aligned and used to build trees (Supplementary Files 2 and 3). For each clade in these two trees, we assessed bootstrap support as high (≥ 50%) or low, depth (mean pairwise tree distance of clade members), and site usage purity (the percentage of clade members using the most frequent integration site).

In graphs of clade depth vs. site purity (Fig. 4) we marked triangles where both axes had low values (i.e., shallow clades with diverse site usage), on the principles of excluding small clades (of four or fewer members) by keeping purity values just below 25% and using the large and diverse clade of promiscuous IS607 (S-Core) integrases to mark the other end of the triangle. The promiscuity triangle thus defined for the S-Int/Core clades was applied directly to the Y-Int clades. These triangles included clades (and their subclades) known for promiscuity: Tn916, Tn4371, tfs and IS607 (red in Supp. Fig. 11 and Supp. Fig. 12). Two known promiscuous clades (CTnDot and ØRSM) were absent, because the number of representatives was too low. Also included in these triangles were two new putative promiscuous clades, one Y-Int and one S-Int. We suspect that the apparent promiscuity in the non-IS607 portions of the S-Core clade have other explanations as described below.

**Fig. 4.**
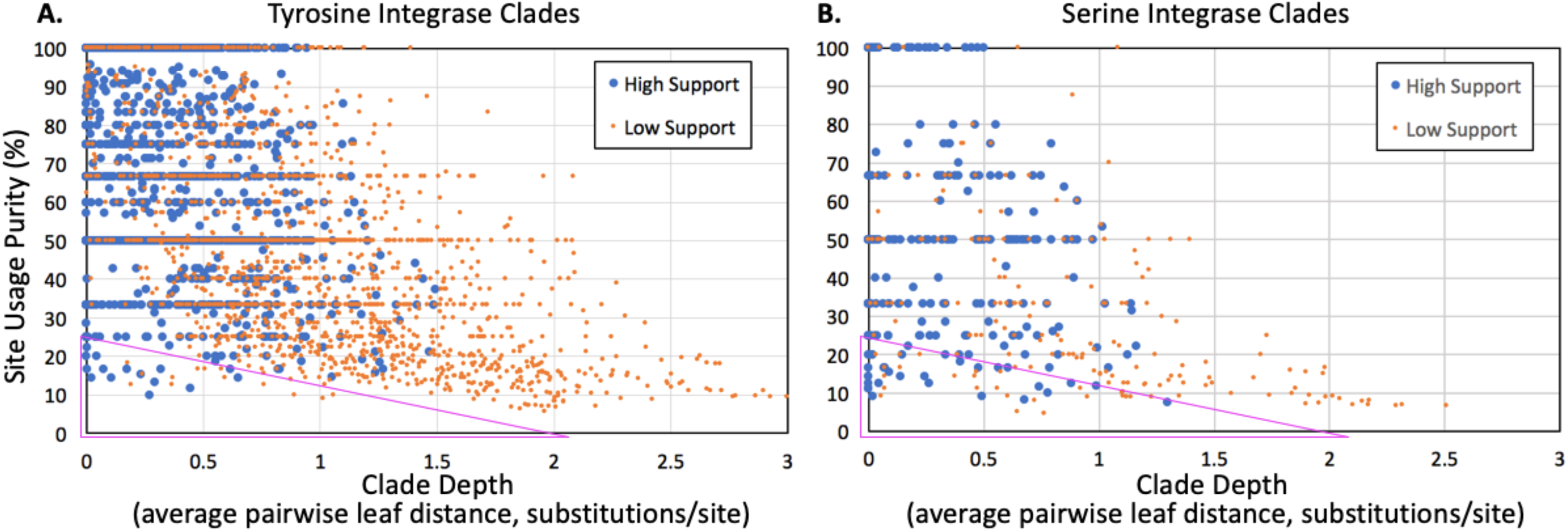
Promiscuous integrase clades. The depth of each clade was measured for each high-support node in the integrase trees (Supp. Files 2 and 3), as was purity of the top site used in the Y-Int (**A**) or S-Int (**B**) clade. The low-depth, low-purity regions (triangles) include known promiscuous clades. Triangles were drawn to include the largest known promiscuous clade (IS607, the furthest right blue dot in the triangle of **B**) and exclude clades of size 4 or smaller.

### Detecting gene inactivation as disruption without restoration

When integrases target protein genes, they typically target an extremity of the gene that is outside of the region encoding conserved protein sequence, or otherwise restore any such conserved coding sequence (CDS) with a similar gene fragment stored in *attP*. That is, such cases need not inactivate the gene; thus, simply detecting that an IGE has invaded a CDS is not sufficient to show that gene function has been inactivated. More rarely, IGEs are known to inactivate genes within conserved peptide-encoding regions, in two situations: 1) accidental targeting of the gene by a promiscuous integrase or by an off-target event from a site-specific integrase, or 2) regulatory gene inactivation as described at *comK, sigK, spsM, mutL, gerE, mlrA, hlb, nifD, fdxN* and *hupL* (4-15). We devised a stringent test for gene inactivation, namely, Pfam domain disruption, by any non-tRNA IGE, comparing top Pfam scores for possible peptides encoded in the *attB* region to those for the *attLR* regions (Table 1). Non-t(m)RNA TIGER IGEs entered domain-coding regions only half as often as mock IGEs from the same genomes (Table 1, line 2), showing a general avoidance of such regions, perhaps in favor of intergenic or gene-terminal *attB*s. Nonetheless we found 410 of the 3308 non-t(m)RNA IGEs disrupting a domain (Supp. File 4). The corresponding fraction for mock IGEs was negligible, showing that determination of gene inactivation constitutes strong additional evidence that the candidate IGE is a true positive. Eight of the 11 cases of known RGI were recovered by this assay (Table 2), although three approached our *attB*:*attLR* bitscore ratio cutoff of 1.1.

**Table 1.** Domain disruption assay. For each mock and non-t(m)RNA TIGER IGE, a 998-bp DNA sequence was taken centered at the left IGE coordinate and likewise at the right coordinate, and all reading frames of each were translated, producing six *attL* and six *attR* peptide sequences. Corresponding IGE-uninterrupted (*attB*) peptides were taken by joining the chromosomal halves of each *attL* and *attR* peptide in all nine positive-frame and nine negative-frame combinations. These 12 *attLR* and 18 *attB* peptide sequences for each IGE were run against the Pfam-A domain HMMs, retaining matches spanning the center crossover position, above a bitscore cutoff of 20 (spanning). For the best-scoring such Pfam for any of the IGE’s *attBLR* peptides, the top score among *attLR* peptides was taken, as was that among *attB* peptides. Occasionally there was no such score for one group (No *attLR*, No *attB*). Otherwise, the top *attB* and *attLR* peptides were compared for that IGE, considering *attB* to “win” and the domain to be disrupted without restoration, if *attB* scored 1.1-fold higher than *attLR*.

|  | TIGER |  | Mock |  |
| --- | --- | --- | --- | --- |
| IGEs | 3308 |  | 32075 |  |
| Spanning (% of IGEs) | 1271 | (38.42%) | 24838 | (77.43%) |
| No *attLR* (% of spanning) | 5 | (0.39%) | 21 | (0.08%) |
| No *attB* (% of spanning) | 28 | (2.20%) | 4230 | (17.03%) |
| *attLR* win (% of spanning) | 828 | (65.15%) | 20521 | (82.62%) |
| *attB* win (% of spanning) | 410 | (32.26%) | 66 | (0.26%) |

**Table 2.** Putative cases of regulated gene integrity. Cases with domain disruption detected are marked in orange; those found as tight integrase clades of size ≤2 are marked in yellow; 5 literature and 19 novel cases had both.

| Broken Pfam | Broken Gene | Integrase Family | Sample Comparator GI(s) | GI Type | Clade Size(s) | Pfam Usage in Clade(s) | Pfam match: attB/attLR | Position in Pfam (%) | Genera | PubMed ref to breakage | Role (quotes: labels commonly found in close homologs' annotations) | Note |
| --- | --- | --- | --- | --- | --- | --- | --- | --- | --- | --- | --- | --- |
| **LITERATURE CASES** |  |  |  |  |  |  |  |  |  |  |  |  |
| Sigma70_r2 | *sigK* | Ser | Bsu14.48.sigK | Prophage | 4 | 3 | 1.183 | 83.1-87.3 | *Bacillus, Anoxybacillus* | 8655528 | Sporulation |  |
| none | *sigK* | Ser | Cdi6.15.sigK | Other | 4 | 4 | N.A. | N.A. | *Clostridioides, Clostridium* | 12694623 | Sporulation | Broken between Sigma70_r2 and Sigma70_r4 Pfam hits |
| Polysacc_synt_2 | *spsM* | Ser | Bam6.20.pseB | Other | 2 | 2 | 1.863 | 42.9 | *Bacillus* | 28535266 | Sporulation |  |
| ComK | *comK* | Ser | Lmo3.40.comK | Prophage | 6 | 6 | 1.585 | 39.5 | *Listeria* | 10652093 | Phagosome escape |  |
| GerE | *gerE* | Ser | Bce7.29.gerE | Other | N.A. | N.A. | 1.340 | 70.7 | *Bacillus* | 28900282 | Sporulation | Omitted from tree due to multiple integrase candidates |
| MerR | *mlrA* | Tyr | Eco282.63.ycgE | Prophage | 5 | 5 | 1.776 | 38.5-42.1 | *Escherichia* | 27190164 | Curli/biofilm production |  |
| Oxidored_nitro | *nifD* | Tyr | Npc1.11.nifD | Other | 1 | 1 | 1.181 | 94.6 | *Nostoc* | 3920531 | Heterocyst | Singleton in tree |
| none | *fdxN* | Ser | not found |  | N.A. | N.A. | N.A. | N.A. | *Nostoc* | 3141375 | Heterocyst | GI not found: 12 reference genomes had q2 hits to Npc1 but with low (76 - 87%) homology and not overlapping the crossover site; multiple *int* candidates; did not break Pfam Fer4_7 |
| NiFeSe_Hases | *hupL* | Tyr | Npc1.9.NiFeSe_Hases | Other | N.A. | N.A. | 1.914 | 38.6 | *Nostoc* | 7846053 | Heterocyst | Omitted from tree due to multiple integrase candidates; not detected as broken because interrupted NiFeSe_Hases segment scored less than a neighboring uninterrupted one |
| Exo_endo_phos | *hlyB* | Tyr | Sau3.42.hlyB | Prophage | 11 | 10 | 1.123 | 5.9 | *Staphylococcus* | 30962356 | β-hemolysin conversion | |
| none | *mutL* | Tyr | Spy1.14.mutL | Other | 9 | 9 | N.A. | N.A. | *Streptococcus* | 18676670 | Mutation rate | Doesn't alter CDS, but separates it from its RBS |
| **NOVEL CASES** |  |  |  |  |  |  |  |  |  |  |  |  |
| AAA_PrkA | *prkA* | Ser | Gym2.15.AAA_PrkA | Other | 2 | 2 | 1.553 | 34.3 | *Geobacillus, Parageobacillus* |  | Sporulation |  |
| AI-2E_transport | *tqsA* | Tyr | Eco33.11.tqsA | Other | 3 | 3 | 1.265 | 10.9 | *Escherichia* |  | Autoinducer-2 export |  |
| DEAD | *hrpB* | Tyr | Ppu4.45.hrpB | Other | 3 | 2 | 1.233 | 75.0 | *Pseudomonas* |  | RNA helicase | GIs contain their own *hrpB* gene |
| DUF1175 | *yfaT* | Tyr | Eco557.50.DUF1175 | Prophage | 2 | 2 | 1.206 | 88.2 | *Escherichia* |  | Unknown |  |
| DUF2466 |  | Tyr | Xau1.104.DUF2466 | ICE | 2 | 2 | 1.663 | 60.5 | *Xanthobacter, Tistrella* |  | "RadC protein used in DNA repair" |  |
| dUTPase | *dut* | Tyr | Rso1.37.dut | Other | 2 | 2 | 1.372 | 53.4 | *Aeromonas, Ralstonia* |  | dUTPase |  |
| FtsK_SpolIIE | *eccCa1* | Ser | Mva2.15.eccCa1 | Other | 2 | 2 | 2.191 | 44.8-45.8 | *Mycobacterium, Rhodococcus* |  | Virulence factor export |  |
| GIDA |  | Ser | Cac3.3.GIDA | Other | 2 | 2 | 2.051 | 21.2-21.3 | *Clostridium* |  | "FAD-dependent Oxidoreductase" | Breaks same gene (JRWL01000001.1/246585-247586) as below |
| GIDA |  | Ser | Cac3.53.GIDA | Prophage | 2 | 2 | 1.159 | 76.7 | *Clostridium* |  | "FAD-dependent Oxidoreductase" | See above |
| GntP_permease | *gntT* | Ser | Lre1.43.gntT | Prophage | 2 | 2 | 1.258 | 11.8 | *Lactobacillus* |  | Gluconate transport |  |
| HNH |  | Ser | Min2.53.DUF222 | Other | 2 | 2 | 1.822 | 71.5 | *Mycobacterium* |  | Homing endonuclease |  |
| HTH_3 |  | Ser | Cdi12.6.immR | ICE | 3 | 2 | 1.697 | 58.2 | *Clostridioides, Enterococcus* |  | Transcription regulation | Not close relative to classical *B. subtilis* ImmR |
| Lon_C | *yjfB* | Ser | Sro2.15.comM, Bca1.12.comM | Other | 3,1 | 2,1 | 1.559 | 53.4-54.0 | *Streptosporangium, Salinispora, Beutenbergia* |  | Mg chelatase-like AAA ATPase | Perhaps erroneously broken into two clades |
| LysE | *rhtB* | Ser | Ppu12.40.rhtB | Other | 10 | 3 | 1.450 | 57.0-57.1 | *Pseudomonas, Burkholderia* |  | Amino acid efflux | Clade also uses similar non-Pfam-breaking *attB* in *rimO* |
| Mg_chelatase | *yjfB* | Ser | Mex5.19.comM | Other | 2 | 2 | 1.168 | 12.4-12.9 | *Nitrobacter, Methylobacterium* |  | Mg chelatase-like AAA ATPase |  |
| Mg_chelatase | *yjfB* | Tyr | Pae17.37.comM | Other | 2 | 2 | 1.335 | 75.2 | *Pseudomonas* |  | Mg chelatase-like AAA ATPase |  |
| Molybdopterin | *ynfE* | Tyr | Eco892.7.ynfE | Other | 2 | 2 | 1.883 | 79.3-79.5 | *Escherichia, Shigella* |  | Selenate/DMS reduction |  |
| Porin_1 | *ompN* | Tyr | Raq2.41.ompN | Prophage | 2 | 2 | 1.912 | 54.6 | *Rahnella* |  | Porin |  |
| Pyr_redox_2 | *merA* | Ser | Sau3.43.merA | Prophage | 3 | 3 | 1.433 | 19.3-20.2 | *Staphylococcus* |  | Redox control |  |
| T4SS-DNA_transf | *traG* | Ser | Cdi9.8.traG | Other | 3 | 3 | 1.863 | 47.7 | *Clostridioides, Streptococcus, Finegoldia* |  | Type 4 secretion system peptidoglycanase | Cross-phylum |

Integrases that thus disrupted domains were marked if present on our integrase trees (as noted above, many had been omitted due to vertical inheritance redundancy or multiplicity of *int* genes in the IGE). In the Y-Int tree, domain disruption occurred for 88 of the 3978 entries, and was significantly enriched in the promiscuous clades Tn916, Tn4371 and *tfs* (8 of their 50 total entries), more than seven-fold the rate for the whole tree, *χ*^2^ (1, *N* = 3978) = 44.5, *p* = 2.5e-11. The rate of domain disruption was much higher in the S-Int tree, at 117 of its 465 entries. Excluding cases accounted for in other ways – the numerous domain disruption cases that we recognized as RGI below (43 instances), the especially large promiscuous IS607 clade (10 instances), and the remainder of the S-Core clade (11 cases) that we cannot convincingly confirm as integrases – we still observe a high rate (53 of 249). This higher rate for S-Ints may be due to unrecognized (singleton) RGI or a mechanistic propensity toward more off-target promiscuity in site-specific clades of S-Ints than for Y-Ints (61,62).

### Regulated gene integrity

We consider a domain disruption case to indicate RGI when found in multiple independent integration events. We found 24 instances where two or more deduplicated IGEs occurred in a tight high-support integrase clade, disrupting the same domain (Table 2). Each was shown to be a unified group sharing the position of disruption within the domain. Despite their unity of site usage, their distributions often crossed genus boundaries and occasionally phylum boundaries.

Five of our detected RGI clades were already known, those affecting *sigK* in *Bacillus, spsM, comK, mlrA*, and *hlb*. Six additional cases known from the literature did not form RGI clades. Three of these (*gerE, nifD*, and *hupL*) did pass our domain disruption test, but did not form clades either because they were the only example in the tree, or had been excluded from the trees because of multiple integrase candidates. For the IGEs inactivating *sigK* in *Clostridium, attB* is not within a Pfam region, but in the space between two. The *fdxN* IGE was not found due to lack of an adequate reference genome. The *mutL*-inactivating IGEs do not affect the coding sequence but separate it from its ribosome binding site. Perhaps with similar functional consequences to the *mutL*-inactivating IGEs, we examined several IGEs in the *mutL* partner gene *mutS* and found their *attB*s at the gene midpoint or further upstream, suggesting inactivation.

For the remaining novel 19 domain-disrupting tight integrase clades, not all the gene functions are known, but they include sporulation, virulence and quorum-sensing functions (Table 2). An additional clade seemed interesting; it included three instances of an inactivated *rhtB* gene but the clade was loosened by its inclusion of seven IGEs integrating (without domain disruption) into the *rimO* rRNA modification gene. The *attB* core sequences are similar at *rimO* and *rhtB*, so it is unclear whether *rhtB* gene inactivation is regulatory or a high-frequency off-target effect of a *rimO*-targeting integrase.

### Gene replacement

In a nuance on RGI, the inactivated (disrupted and unrestored) target gene may be supplanted by an IGE-borne homolog of that same gene, as reported for a tmRNA-encoding IGE integrated into a *Rhodobacter* tmRNA gene (63). We note here that the *hrpB*-inactivating islands encode a different *hrpB* gene. Such replacement may simply act to restore the function of the inactivated target gene or may shift phenotype if the replacement has a slightly altered function.

## DISCUSSION

Regulated gene integrity and integrase promiscuity were approached by software for comprehensive and precise mapping of IGEs in archaeal and bacterial genomes, combined with integrase phylogenetic analysis and a gene inactivation test. We found that RGI is more widespread than the literature had indicated, and discovered several new candidate cases, evidenced by identifying multiple independently IGE-interrupted and multiple uninterrupted versions of each regulated gene. The implied switching of gene activity by IGE integration and excision already meets a minimal definition of regulation, and these cases may rise to a stronger significance of regulation, where switching is linked to a particular physiological situation that benefits fitness of either the host or the IGE. To confirm this will require sufficiently deep understanding of the physiological roles of the affected genes among their various bacterial hosts to identify experimental manipulations that induce regulatory excision or integration.

Although many cases in our list of domain-disrupting IGEs (Supp. File 4) arose through promiscuity, we expect that additional cases of RGI are also among them, missed perhaps because they were single-leaf clades or not analyzed in our trees; candidates are those disrupting Pfams GreA_GreB_N and IstB_IS21. Application of our methods to the ∼300,000 bacterial genomes now available will certainly add more. Furthermore, our domain disruption assay was too stringent to catch all classical cases of RGI. Future improvements will call sites *between* an incontrovertible start codon and a Pfam match, include in trees all integrases from multi-*int* IGEs, and simply exclude the confounding S-Core proteins. Relaxing the stringency of our gene inactivation test is expected to yield more new cases; the four *int* clades we found targeting *mutS* are candidates, given that its partner gene *mutL* is already a known target of RGI (13); inspection shows that these cases disrupt the genes upstream of MutS Pfam matches. There may even be ways to go beyond Pfams to discover RGI in hypothetical genes.

The S-Core proteins have traditionally been known for resolvase and DNA invertase functions. An exception is that from ØRSM, which has been shown to function as a site-specific integrase (22); in our tree this integrase had only one reasonably close relative and therefore could not reveal its site-specificity. Indeed the S-Core group shows no evidence of site-specificity in our tree. The large IS607 S-Core clade was used to define the promiscuity zone of Fig 4B and some additional clades in the S-Core group do fall within the zone (Supp. Fig. 12), but many do not because they appear as long isolated branches. The group is underrepresented among the IGE calls (Fig 3C), and those that were called may be explained by non-integrase resolvase and invertase functions as follows. Some calls may be from transposable elements encoding S-Core resolvases, as at Sau15.7.trmB|ytnP; S-Core IGE calls contain transposase genes at higher rates than other integrase groups (32.2% S-Int, 30.2% Y-Int and 25.8% S-Core IGE calls), and they more often fall in the short length range of ISs (67.8% S-Core, 35.1% S-Int and 19.7% Y-Int IGE calls are < 10 kbp), as might be expected for resolvases in ISs/transposons. DNA invertases tend to control expression of genes that may be subject to negative selection; thus these S-Core genes may be subject to deletion by processes other than action of the S-Core itself. The group also shows the lowest support values (35.3% S-Core, 17.9% S-Int, and 14.7% Y-Int IGE calls score < 5% of the maximum support value for the genome). For these reasons we suspect that, apart from the ϕRSM and IS607 clades, the S-Core protein family contains few IGE integrases.

The TIGER program is based on principles of cohesion of the IGE integration module and comparative genomics, implemented through ping-pong BLAST. We suspect that, to manually map IGEs, other researchers have intuitively used an approach similar to that automated in TIGER. Merging with our previous software Islander, which is limited to IGEs in t(m)RNA gene targets, improves yields. Three approaches were taken to ruling out false positive IGE calls: rejection of query-reference genome matches based only on matches to transposons, a formula combining multiple IGE metrics, and identification of regions of frequent genomic rearrangement due to agents other than the candidate integrase. Except for missing a few IGEs split into separate scaffolds, preliminary testing has shown that TIGER (and Islander) performs nearly as well on incomplete genomes as on complete genomes. We have also found that few reference genomes are from a different taxonomic class than the query genome; using smaller class-based reference genome databases will speed future scale-up. The precise mapping of IGEs is an advancement in bioinformatics, with practical benefits. For example, in our phage therapy work, it allows phage genome engineering design to begin immediately.

The frequent occurrence of multiple integrase genes in the same IGE serves as a cautionary note that even when an IGE contains only one *int* gene, its site-specificity may not be mapped; the *int* responsible for an IGE’s location may have been deleted from a multi-*int* IGE.

Certain categories of IGEs were missed with this first instance of TIGER. These include IGEs such as phage Mu that encode integrases from other protein families, GIs such as CTXϕ that do not encode their own integrases but exploit housekeeping enzymes (64), and GIs whose integrase gene has degraded beyond recognition; the former problem is solved by allowing integrases from other families. We consider Islander and TIGER to be subject to different types of false positives. Islander artifacts tend to arise from casual matches to tRNA gene BLAST queries, and therefore tend to be random DNA segments. Barring assembly errors, TIGER mainly finds real insertion/deletion relationships between genomes. These are usually due to normal action of the integrase gene on its IGE, but may occasionally arise from other recombinational processes that do not properly map an IGE, as movement of IGE-surrounding integron cassettes may do.

It has long been asked why IGE integrases preferentially target tRNA gene sites (58). A “symmetry hypothesis” is attractive, in which this bias comes from the preference of Y-Ints for *attB* site dyad symmetry which neatly matches the inverted repeat symmetry and spacing found at all t(m)RNA gene anticodon and TΨC stem-loop segments. However fine mapping of Y-Int t(m)RNA gene *attB*s showed that approximately half are instead at an asymmetrical site between the TΨC and accteptor stems, and therefore not explainable by the symmetry hypothesis (59). If the preference for t(m)RNA was based solely on the evolutionary rationale that their primary sequences are conserved, allowing more reliable *attB* presence in new hosts, the two integrase families should prefer t(m)RNA genes at similar frequencies. Instead, we found that Y-Ints target t(m)RNA genes at a significantly higher rate than do S-Ints. Perhaps during evolution of new site-specificity, some other mechanistic aspect of Y-Int action (beside symmetry preference) tends to direct them specifically toward t(m)RNA genes, and this mechanism does not pertain to S-Ints.

