## Supplementary material for "New candidates for regulated gene integrity revealed through precise mapping of integrative genetic elements": Supp. Fig. 1-12

SUPPLEMENTARY MATERIALS
SUPPLEMENTARY TABLES (See separate files for Supp. Tables 1-4, 10 and 11)

Supp. Table 5. Validation of IS discovery by TIGER in the *E. coli* MG1655 genome. The well-annotated genome of *E. coli* MG1655 (NC_000913.2) has 47 IS calls that were identified with ISFinder entries where possible, and sorted into three categories: 1) 26 intact with the expected direct repeat (DR) at the flanks, 2) 13 intact without the expected DR, and 3) 8 truncated. The latter two categories may not be the immediate products of transposition but have more complicated evolutionary histories, in category 2 because they may be products of homologous recombination between two copies of the same IS.

TIGER was applied in IS mode at all 79 transposase genes detected in the genome, yielding 35 IS calls and leaving only six transposase genes unassigned. TIGER calls were matched or overlapped to the standards as possible. None of the category 3 (truncated) and only two of the category 2 (no-DR) standards were matched by TIGER calls; these categories were less likely to preserve the original post-transposition flanking sequences as required by TIGER. For the 26 category 1 (DR-flanked) IS standards, 22 were identified accurately, with precise or nearly precise (subject to blast hit terminus drift) identification of the DR; the four that were missed could be ascribed to nearby recombination events that truncated the original flanking regions to shorter than our 500 bp minimum. The 11 TIGER calls not matching standards (Overlapping but not matching, and TIGER-only) probably reflect chromosomal deletions in reference genomes by rare mechanisms other than transposition; they had very low support values, suggesting that these false positives can be controlled.

The ΔL column is the distance between the left TIGER coordinate (centered within the DR (direct repeat) segment) relative to the IS coordinate, likewise for ΔR at the right. ExpDR is expected DR length from ISFinder. The DR column shows sequences at the left (above) and right (below) for cases where DRs were identified (in red); lower case shows the DR sequences defined by TIGER and IS sequence according to ISFinder is underlined.

| Gen-Bank Name | ISFin-der Name | Left | Right | TIGER Support | ΔL | ΔR | Exp DR | DR |
|---|---|---|---|---|---|---|---|---|
| **MATCHING (24)** | | | | | | | | |
| IS3E | ISEc17 | 2168195 | 2169452 | 9228 | 2 | -2 | ? | GAGAATAACGgtcTGATCTTACC…<br>…GGTAGGATCAgtcATCCCTTCGT |
| IS186C | IS186A | 2512295 | 2513639 | 8893 | 3 | -3 | 10 | AAAAAAGCCGgggataattcccATAAGCGCTA…<br>…TAGCGCTTATgggataattcccCGGTTTTTTT |
| IS5F | IS5 | 1394068 | 1395262 | 8551 | 3 | -3 | 4 | GCAACATTAGctaaGGAAGGTGCG…<br>…CGCACCTTCCctaaCTAATCAATG |
| IS1H | IS1A | 1976527 | 1977294 | 8436 | 5 | -5 | 9 | AATTACTTAAcataaatgGGTAATGACT…<br>…CAGCATCACCcataaatgTATAAGTCAT |
| IS2A | IS2 | 380484 | 381814 | 7458 | 4 | -3 | 5 | ATAAACGCATaattacTAGACTGGCC…<br>…GGGCAAATCCaattacCTATCAGGCA |
| IS5K | IS5 | 2286941 | 2288135 | 6235 | 3 | -3 | 4 | ATGGGTGAATataaGGAAGGTGCG…<br>…CGCACCTTCCataaCGCTGTAGCC |

| | | | | | | | | |
|---|---|---|---|---|---|---|---|---|
| IS1D | IS1D | 1049001 | 1049768 | 6081 | 5 | -5 | 9 | TCATCGCATGgacaatacgGGTGATGCTG…<br>…AGTCATTACCgacaatacgTACGCTTGAG |
| IS1E | IS1A | 3581451 | 3582218 | 5819 | 5 | -5 | 9 | ATTTACTTATgacattaaaGGTGATGCTG…<br>…AGTCATTACCgacattaaaAGTAACTTTT |
| IS5T | IS5 | 3650059 | 3651253 | 5315 | 2 | -3 | 4 | TGACGCAAATttaggGAAGGTGCGA…<br>…CGCACCTTCCttaggTGACACTATT |
| IS2I | IS2 | 3184118 | 3185448 | 4792 | 6 | 1 | 5 | ATTGTTCATTatcc-gttATTAGACTGG…<br>…ATAGGGGCAAatccagtTATCATCATTG |
| IS186A | IS186A | 15387 | 16731 | 4628 | 3 | -3 | 10 | TAAAGTGGCGgggatcactcccATAAGCGCTA…<br>…TAGCGCTTATgggatcactcccCGCCGTTGCT |
| IS5H | IS5 | 2064183 | 2065377 | 4392 | 3 | -3 | 4 | CGACTATGCActagGGAAGGTGCG…<br>…CGCACCTTCCctagAACACCACAA |
| ISZ | none | 1293649 | 1294628 | 3728 | 15 | -6 | ? | AAAATACGGCcccagaagggCAATGCCGTT…<br>…ATCGGCATTGcccagaagggGCCGTTTATG |
| IS5D | IS5 | 687074 | 688268 | 3011 | 2 | -3 | 4 | GTTGTGCCTGttaggGAAGGTGCGA…<br>…CGCACCTTCCttaggTAACATTTAG |
| IS186B | IS186B | 607230 | 608574 | 2319 | 3 | -3 | 8 | CCGTTGTGCGgggagtaatcccATAAGCGCTA<br>…TAGCGCTTATgggagtaatcccCGCATATCCG |
| IS3B | IS3 | 390933 | 392190 | 1370 | 2 | -3 | 3 | GTGGGGGGCTgattGATCCTACCC…<br>…GGTAAGATCAgattACAACCTGTA |
| IS5B | IS5B | 573814 | 575008 | 1151 | 0 | -5 | 4 | ACCGAGTGCTttagggaagGTGCGAATAA…<br>…CGCACCTTCCttagtgaagTCATTTTTGT |
| IS2K | IS2 | 4496204 | 4497534 | 1028 | 3 | -4 | 5 | TTTGATGGTCccttgtAGACTGGCCC…<br>…GGCAAATCCAccttgtGCTGATATGA |
| IS5R | IS5 | 3363578 | 3364772 | 831 | 3 | -3 | 4 | ACAGCATCAAttagGGAAGGTGCG…<br>…CGCACCTTCCttagTACTTCCCCC |
| IS4 | IS4 | 4500113 | 4501538 | 241 | 8 | -5 | 11 | CGAAATAGGCatcttggttctgTAATGCCAGT…<br>…ATCGGCATTAatcttggttctgGTGTTTGTAA |
| IS5I | IS5 | 2099773 | 2100967 | 82 | 3 | -3 | 4 | TTTTGTTTTAttaaGGAAGGTGC…<br>…CGCACCTTCCttaaTCCCTAACA |
| IS3D | IS3 | 1093468 | 1094725 | 78 | 6 | -4 | 3 | No DR detected |
| IS150 | IS150 | 3718656 | 3720098 | 64 | 7 | -2 | 3 | No DR detected |
| IS2D | IS2 | 1465934 | 1467264 | 5 | 3 | -3 | 5 | ATGGGACAAGaacccTGGATTTGCC…<br>…GGCCAGTCTAaacccCAGAACTTAC |
| **OVERLAPPING BUT NOT MATCHING (7)** | | | | | | | | |
| IS3C | IS3 | 566000 | 567257 | 9 | 950 | -6555 | 3 | No DR detected |
| IS600 | IS600 | 4506703 | 4507029 | 8 | 148 | -722 | 3 | Partial IS |
| IS2E | IS2 | 1648867 | 1649572 | 7 | 1814 | -1290 | 5 | Partial IS |
| IS2H | IS2 | 2994383 | 2995713 | 3 | 8278 | -608 | 5 | No DR detected |
| IS911A | IS911 | 269430 | 269764 | 2 | -749 | -1080 | 3 | Partial IS |
| IS1B | IS1B | 278387 | 279154 | 1 | 754 | -10703 | 9? | No DR detected |

| | | | | | | | | |
|---|---|---|---|---|---|---|---|---|
| IS2F | IS2 | 2066965 | 2068295 | 1 | 1586 | -8111 | 5 | ATCCGCATTCGTGGCTGGATTTGCC…<br>…GGCCAGTCTAGTGGCGAAGCATCCT |
| **MISSED BY TIGER (16)** | | | | | | | | |
| IS1A | IS1A | 19796 | 20563 | | | | 9 | No DR detected |
| IS30A | IS30 | 269765 | 270985 | | | | 2 | AAAAAAGGCTACTGTAGATTCA…<br>…TGAATCTACAACCGCGCTCTTG |
| IS911A | IS911 | 270986 | 271414 | | | | 3 | Partial IS |
| IS5A | IS5 | 273179 | 274373 | | | | 4 | No DR detected |
| IS30B | IS30 | 279155 | 279335 | | | | 2 | Partial IS |
| ISX | ISEhe3 | 279338 | 279649 | | | | 3 | Partial IS |
| IS1C | IS1B | 289858 | 290625 | | | | 9? | No DR detected |
| IS3A | IS3 | 314450 | 315707 | | | | 3 | No DR detected |
| IS5Y | IS5 | 1425623 | 1426818 | | | | 4 | TAACTGACAACAAGGGAAGGTGCG…<br>…CGCACCTTCCCAAGGGGAAAACGC |
| IS30C | IS30 | 1467320 | 1468540 | | | | 2 | AATAAAGGTACTTGTAGATTCA…<br>…TGAATCTACACTGTTGAAATTC |
| IS609 | IS609 | 1501185 | 1502932 | | | | ? | No DR detected |
| IS5LO | IS5D | 3128168 | 3129362 | | | | 4 | No DR detected |
| IS911B | IS911 | 4505148 | 4505481 | | | | 3 | Partial IS |
| IS30D | IS30D | 4505482 | 4506702 | | | | 2 | No DR detected |
| IS911B | IS911 | 4507030 | 4507824 | | | | 3 | Partial IS |
| IS1F | IS1F | 4516495 | 4517262 | | | | 9? | No DR detected |
| **TIGER-ONLY (4)** | | | | | | | | |
| | | 736800 | 738067 | 17 | | | | |
| | | 4566855 | 4577803 | 12 | | | | |
| | | 1525914 | 1531649 | 1 | | | | |
| | | 3617011 | 3623698 | 1 | | | | |

Supp. Table 6. Evaluation of TIGER stages. Raw calls from tigercore.pl were merged (TIGER raw). After running resolve.pl, calls identified as false positives by our formula (Supp. Fig. 4) were included (TIGER + FP) or excluded (TIGER final). These datasets were evaluated as GIs (loose definition) using the 104-genome system of Bertelli et al, 2019 (Brief Bioinform **20**:1685-1698). Here "precision" indicates lack of overlap to GI-negative chromosomal regions of the test system.

| IGE set | Total length (Mbp) | Precision | Recall | F-score |
|---|---|---|---|---|
| TIGER raw | 31.7 | 0.950 | 0.438 | 0.511 |
| TIGER+FP | 21.3 | 0.996 | 0.404 | 0.497 |
| TIGER final | 20.0 | 1.000 | 0.396 | 0.489 |

Supp. Table 7. *Vibrio* IGEs masked by superintegron activity. IGEs are grouped in three integrase clades. Coordinates are midpoints of flanking *attC*s.

<u>S-Int clade</u>
NC_009456.1:813973-832951 (PLE5; *V. cholerae* 0395)
NC_012583.1:430991-449969 (PLE5; *V. cholerae* 0395; independent assembly)
NC_012667.1:939186-958281 (PLE4; *V. cholerae* MJ-1236)

<u>Y-Int clade 1</u>
NC_004459.3:2490421-2510912 (*V. vulnificus* CMCP6)
NC_005139.1:1807656-1839081 (*V. vulnificus* YJ016)
NC_013456.1:1840317-1855147 (*V. antiquarius* EX25)

<u>Y-Int clade 2</u>
NC_013456.1:1779422-1825521 (V. *antiquarius* EX25, in same superintegron as above)

Supp. Table 8. Top 95 *attB* sites. Sites are named according to the gene(s) they are in or flanking. t(m)RNA genes are given in one-letter code for the charging amino acid (Z, tmRNA; B, initiator tRNA). Other RNA gene names are those from Rfam. For protein genes, preference is for 1) mnemonic gene name as given by Prokka, 2) top Pfam hit, 3) protein product name from Prokka, otherwise 4) it is designated HYP (hypothetical). Pipe symbol, intergenic space between the flanking gene names; plus symbol, a region spanning genes. PubMed IDs for references to usage of non-t(m)RNA sites are given in red, with named example prophages in parentheses.

| Count | Target, PubMedID (Example) |
|---|---|
| 476 | R |
| 439 | L |
| 412 | S |
| 291 | T |
| 226 | Z |
| 189 | F |
| 171 | HYP |
| 165 | P |
| 155 | K |
| 153 | G |
| 149 | V |
| 134 | I |
| 112 | M |
| 108 | N |
| 94 | A |
| 77 | guaA, 24372381 (ICELm1) |
| 65 | E |
| 60 | U |
| 56 | Y-Int |
| 51 | rlmCD, 26779141 |
| 47 | B |
| 45 | Q |
| 42 | Y |
| 40 | HYP\|HYP |
| 39 | C |
| 27 | proP, 16040988 (Kim) |
| 26 | ychF, PMC5363835 (ST556-0.03) |
| 25 | D |
| 25 | YicC_N, 20807202 (MGIVvuTai1) |
| 24 | rlmH, 29079617 (ψSCCmec) |
| 24 | W |
| 23 | DUF2466 |
| 22 | hlb, 8454180 (Sa3int) |
| 22 | ssb |
| 21 | fimD |
| 21 | ttcA, 26530864 (ϕRac) |
| 20 | rplL, 24372381 (ICESsuSC84) |
| 19 | sufB, 28348852 (ϕSa1) |
| 19 | tcrA |
| 19 | thiL |
| 18 | groL |
| 18 | mnmE, 15752209 (SGI1) |
| 18 | potB, 17351047 (ϕ-CFT073-potB) |
| 18 | prfC, 18326579 (ICEPdaSpa1) |
| 18 | RybB_RNA |
| 17 | FliB, 12142430 |
| 17 | LamB_YcsF\|nei |
| 17 | mutS, 18676670 (SpyCIM1) |
| 17 | ompW, 22135463 (cpO) |
| 16 | pepN\|pncB, 10411733 (Gifsy-2) |
| 15 | HYP\|betA |
| 14 | rpsD, 25161960 (SagCI) |
| 13 | 6S_RNA |
| 13 | CyaR_RyeE_RNA |
| 13 | hup |
| 13 | rplS |
| 12 | lepA, 10411733 (Gifsy-1) |
| 12 | metQ, 30873127 (SaPI1) |
| 12 | sbcB,18824528 (ϕ2851) |
| 12 | STnc540_RNA |
| 11 | DUF222 |
| 11 | GIDA |
| 11 | mutL, 18676670 (ϕSF370.4) |
| 11 | wrbA, 10734605 (ϕσπ5) |
| 11 | yfhL, 18757547 |
| 11 | YolD |
| 10 | DUF177\|DUF795 |
| 10 | GSPII_E |
| 10 | rpmE2 |
| 9 | comK, 10652093 (ϕA118) |
| 9 | cpxP\|fieF |
| 9 | DUF2545 |
| 9 | gor |
| 9 | lip2 |
| 8 | DUF72 |
| 8 | HYP+L |
| 8 | pflA |

Supp Table 9. Benchmarking analysis. All data are from Bertelli et al, 2019 (Brief Bioinform **20**:1685-1698), except for the line for TIGER, which was newly run using the software and test GI (loose definition) data provided. Predictors are sorted by 104-genome precision, which indicates lack of overlap to GI-negative chromosomal regions of the test system.

| | 104-genome GI-positives/negatives | | | 6-genome gold standard GIs | | |
|---|---|---|---|---|---|---|
| Predictors | Precision | Recall | F-score | Precision | Recall | F-score |
| TIGER | 1.000 | 0.396 | 0.489 | 1.000 | 0.599 | 0.728 |
| Islander | 0.971 | 0.140 | 0.204 | 1.000 | 0.226 | 0.354 |
| VRprofile | 0.940 | 0.417 | 0.511 | 0.993 | 0.542 | 0.643 |
| GIHunter | 0.934 | 0.634 | 0.713 | 0.981 | 0.745 | 0.832 |
| SIGI-CRF | 0.920 | 0.400 | 0.498 | 0.993 | 0.436 | 0.522 |
| SIGI-HMM | 0.919 | 0.264 | 0.374 | 0.998 | 0.285 | 0.420 |
| IslandViewer 4 | 0.904 | 0.732 | 0.780 | 0.998 | 0.619 | 0.753 |
| IslandPath-DIMOB | 0.874 | 0.469 | 0.554 | 0.979 | 0.521 | 0.669 |
| MSGIP | 0.865 | 0.163 | 0.199 | 0.947 | 0.353 | 0.439 |
| Zisland Explorer | 0.853 | 0.177 | 0.239 | 0.833 | 0.171 | 0.264 |
| PAI-IDA | 0.685 | 0.149 | 0.220 | 0.667 | 0.067 | 0.118 |
| INDeGenIUS | 0.651 | 0.275 | 0.356 | 0.716 | 0.189 | 0.293 |
| GC-Profile | 0.637 | 0.225 | 0.205 | 0.667 | 0.034 | 0.063 |
| Centroid | 0.627 | 0.217 | 0.292 | 0.636 | 0.129 | 0.202 |
| GI-SVM | 0.603 | 0.317 | 0.382 | 0.721 | 0.200 | 0.312 |
| AlienHunter | 0.594 | 0.570 | 0.540 | 0.753 | 0.570 | 0.642 |
| PredictBias | 0.577 | 0.623 | 0.526 | 0.868 | 0.817 | 0.838 |
| Wn-SVM | 0.552 | 0.286 | 0.354 | 0.715 | 0.208 | 0.320 |
| MTGIpick | 0.551 | 0.675 | 0.559 | 0.819 | 0.744 | 0.775 |
| MJSD | 0.521 | 0.638 | 0.497 | 0.742 | 0.697 | 0.694 |
| SigHunt | 0.457 | 0.605 | 0.466 | 0.692 | 0.624 | 0.649 |

SUPPLEMENTARY FIGURES

Supp. Fig. 1. Definitions of classes of mobile DNA elements. Parentheses denote the associated transmission vehicle (bacteriophage particles or conjugation pili).

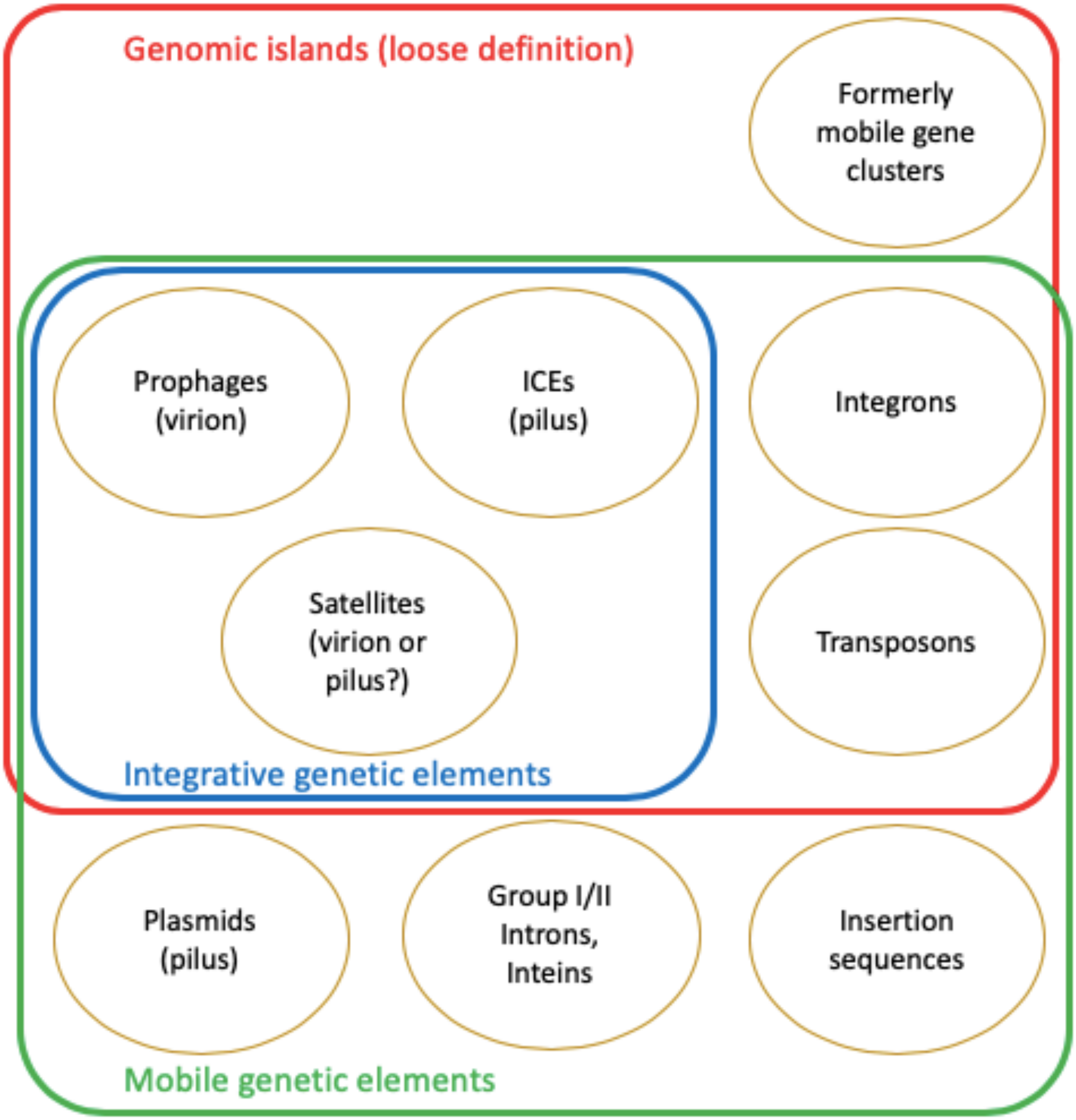

Supp. Fig. 2. IGE integration within a gene, without inactivating the gene. For an IGE targeting a t(m)RNA gene, a fragment of that gene present in *attP* take the place of the equivalent fragment displaced by integration.

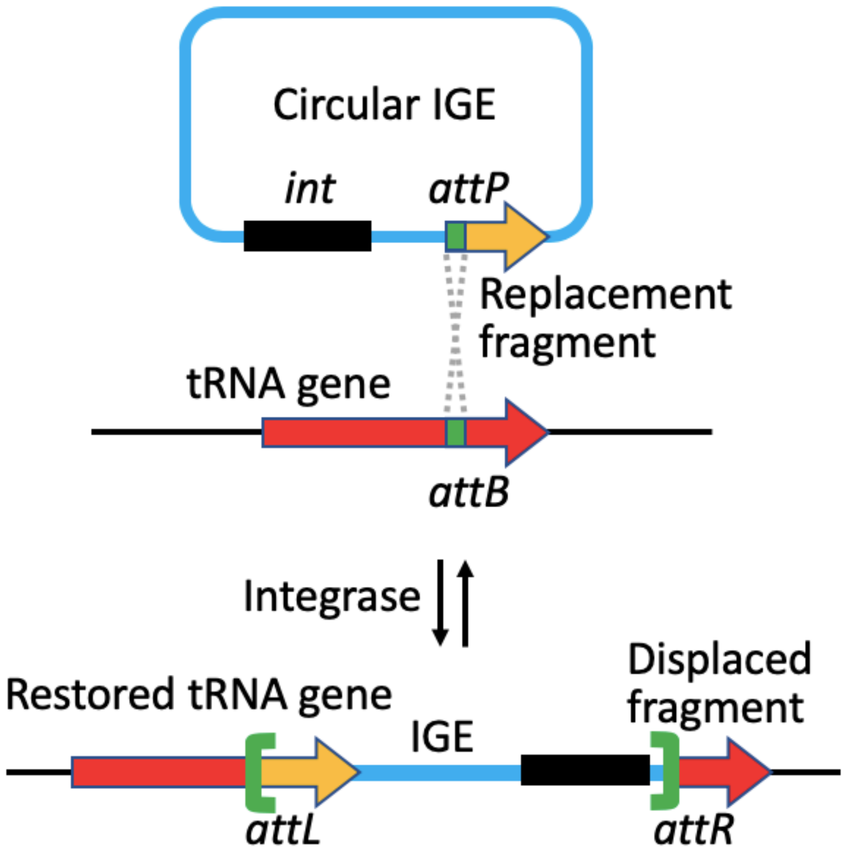

Supp. Fig. 3. Score distributions for seven IGE metrics. Each set of scores was scaled, and length, delta Int and absolute value of mononucleotide bias were log10 transformed. Red, the full-negative IGEs that appeared in an outer quartile (dashed line) for all metrics.

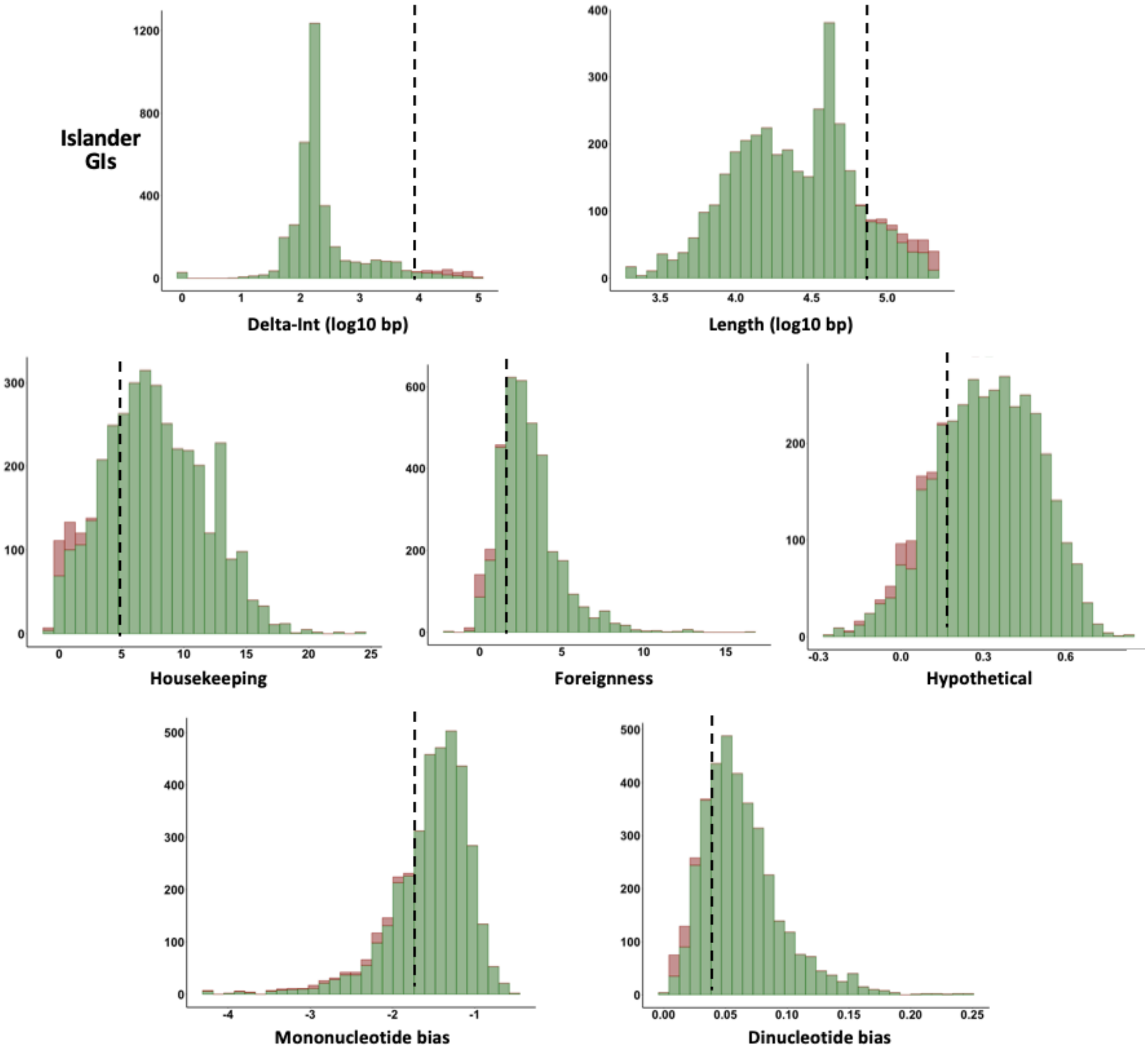

Supp. Fig. 4. Development of a false positive formula. **A**) Principal component analysis of Islander IGEs as in Fig 1A but with doubly-confirmed IGEs marked. **B**) Convex hull drawn around the TIGER-confirmed islands, establishing a "true positive zone" of doubly-confirmed IGEs. **C**) Re-categorization of islands based on their position with respect to the hull. **D**) Formula derived by linear regression on these two new categories of the IGEs. **E**) Final categorization into true and false positives based on the formula.

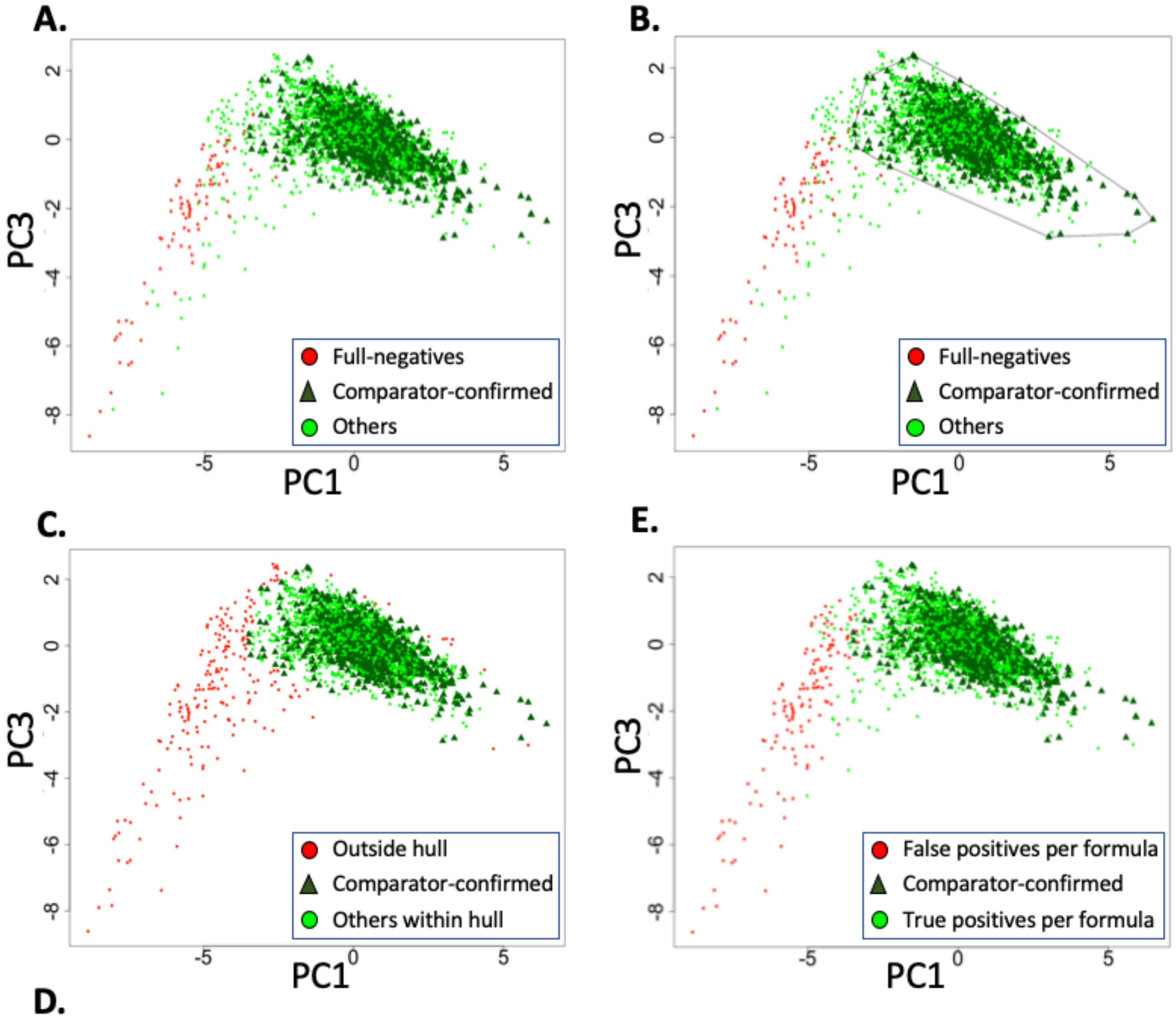


D.

$$\text{FPscore} = \exp(30.6 + -8.3*\log10(\text{length}) - 0.33*\log10(\text{dInt}) - 3.21*(\text{foreign}) - 1.71*(\text{housekeeping}) - 22.3*(\text{hypothetical}) - 16.0*(\text{deltaGC}) - 149.2*(\text{dinucl}))\ [\text{cutoff} = 0.51]$$

Supp. Fig. 5. Genomic instability masking any that may be due to the IGE PLE5 *int*. The integrase of the *Vibrio cholerae* O395 superintegron rearranges the integron by moving cassettes at the *attC* sites (marked in red), masking the mobility signal from the PLE5 IGE within. PLE5 integration split an *attC* (green X's). Raw TIGER calls are shown by bars with support values.

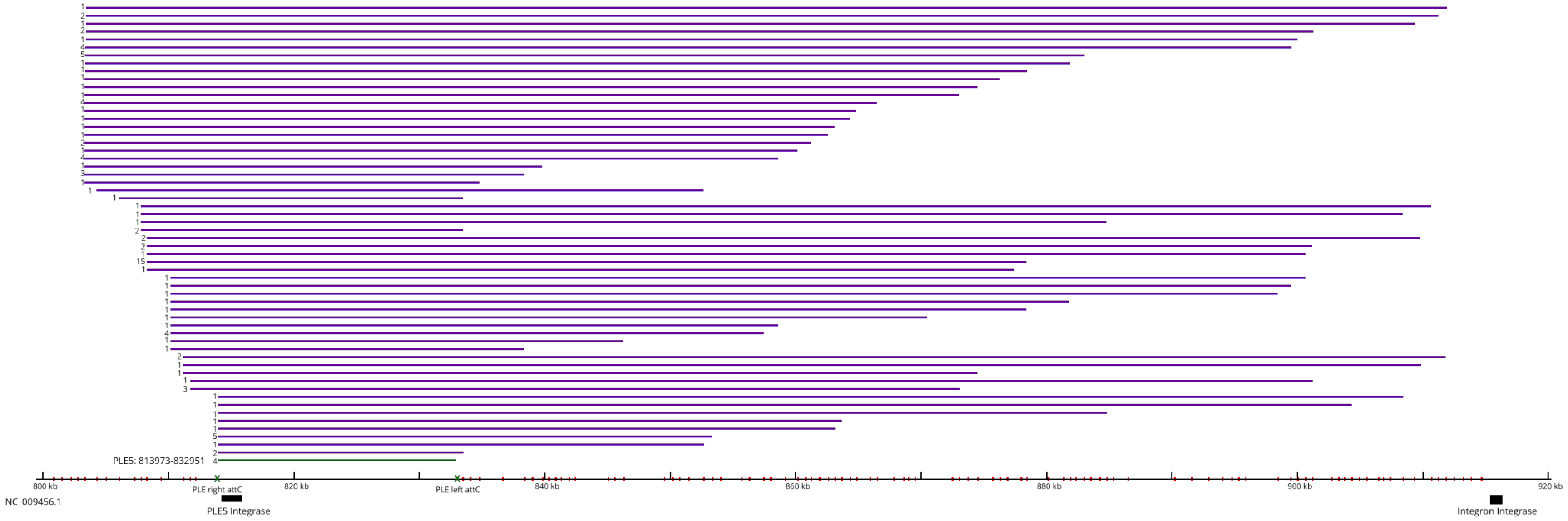

Supp. Fig. 6. Unstable genomic regions. Top:second support ratio was taken for each of 2510 *int*s with multiple TIGER IGE calls. Integrase genes falling below the solid black curve, including the circled zone for *Vibrio* superintegrons, and any IGE calls that depended on them, were rejected as being in regions whose instability is less well ascribed to the *int* itself.

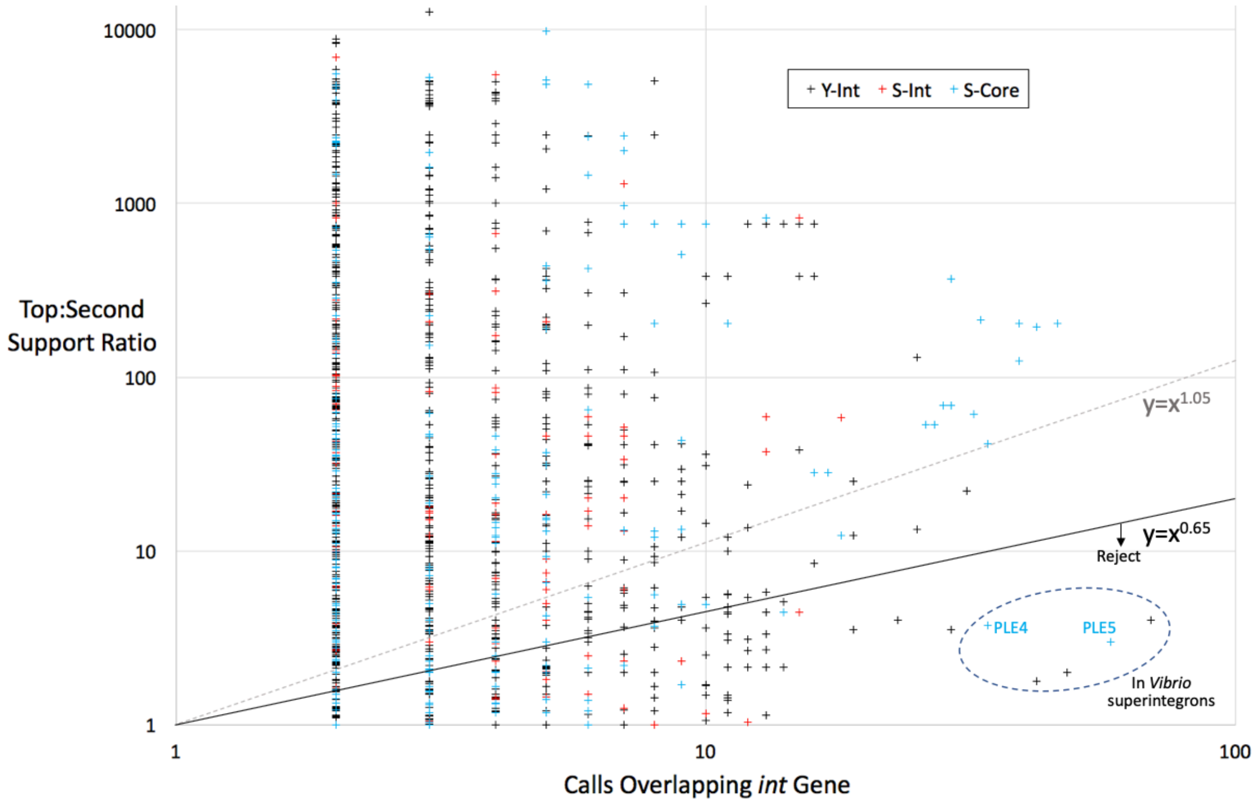

Supp. Fig. 7. Cutoff curve for the top:second support ratio plot. Power-law curves $\mathbf{y = ax^k}$ were drawn through Supp. Fig. 3, where **a**=1 (blue) or **a**=5 (orange) and 0.05 increments of **k**. Effects on rejection rates of all IGEs, Islander-confirmed IGEs, and IGEs with only S-Core integrases were considered; rejection rates at discontinuities in these plots suggested the cutoff curve $y = x^{0.65}$.

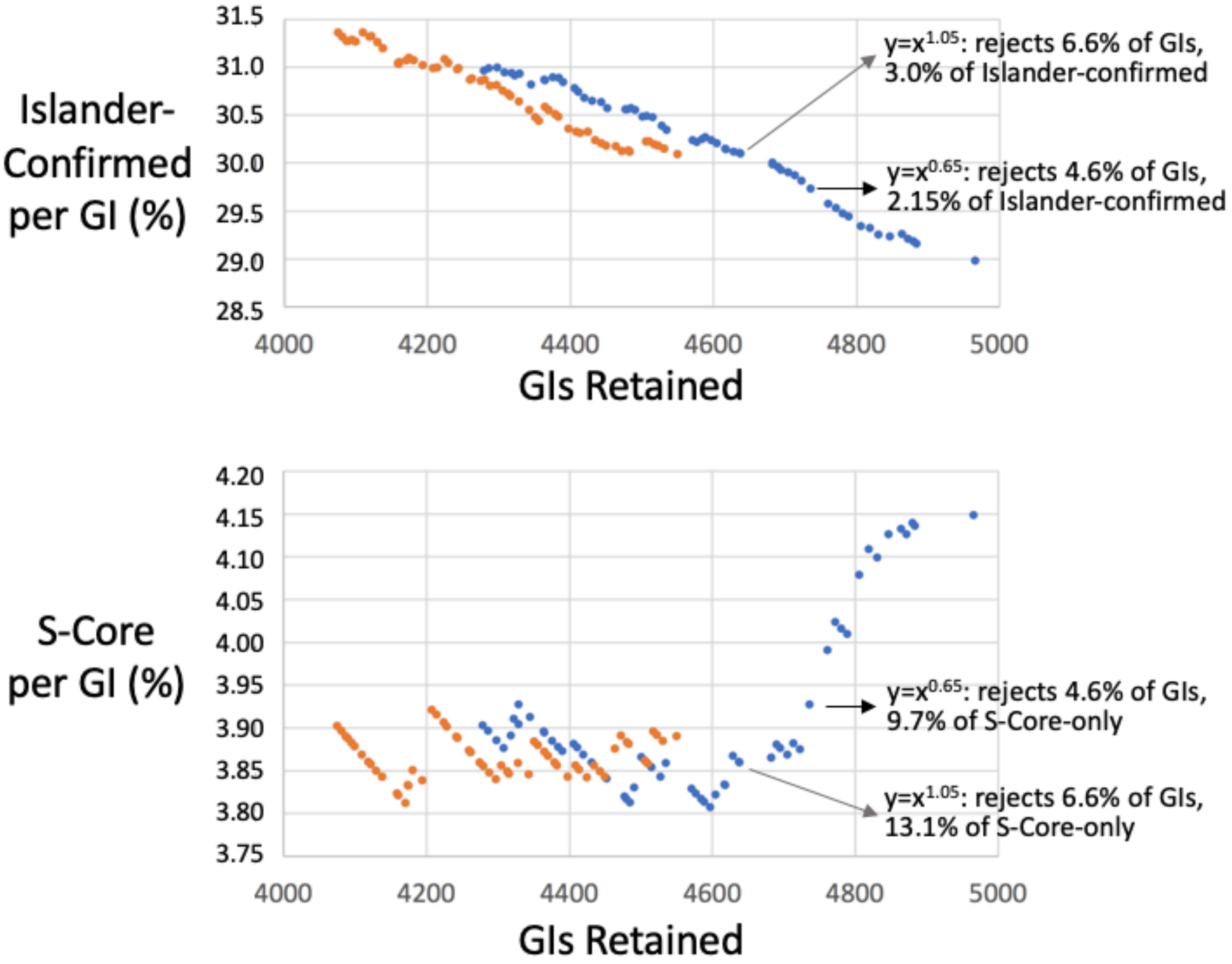

Supp. Fig. 8. Taxonomic distribution of IGEs. Average IGE/genome values are shown by phylum, or by class for the two largest phyla. Genome counts in parentheses.

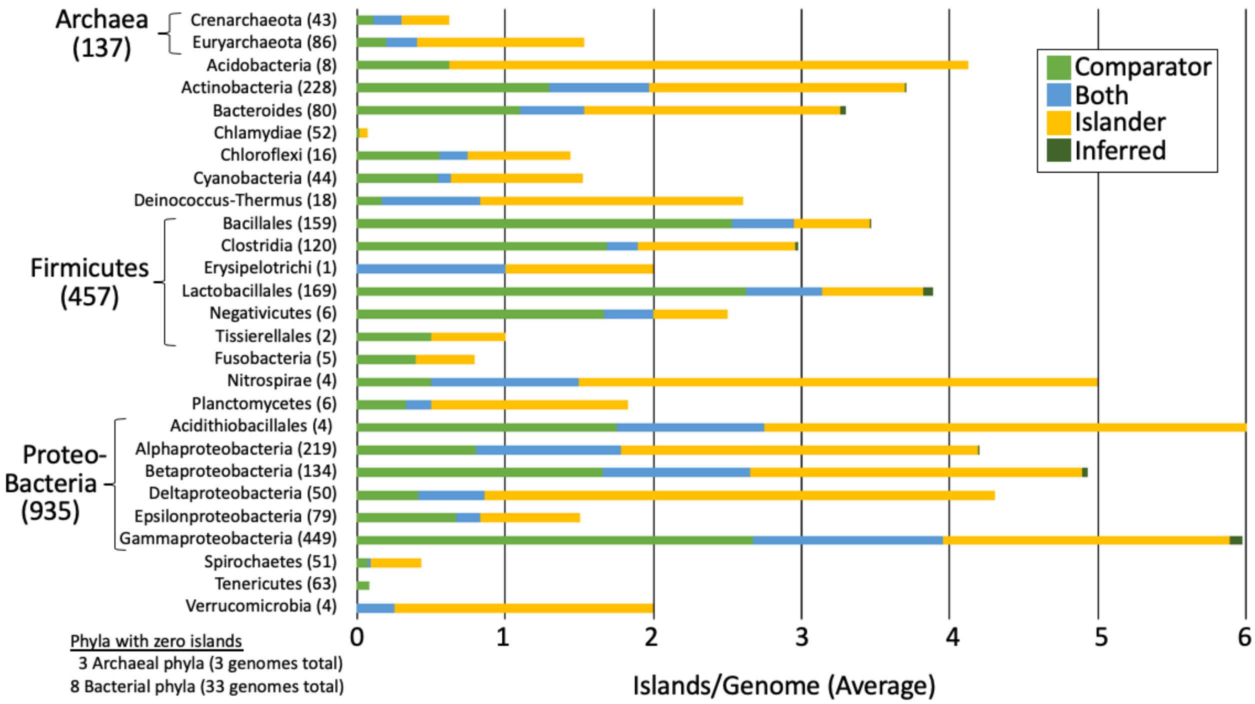

Supp. Fig. 9. Support values as a function of phylogenetically close genome counts.

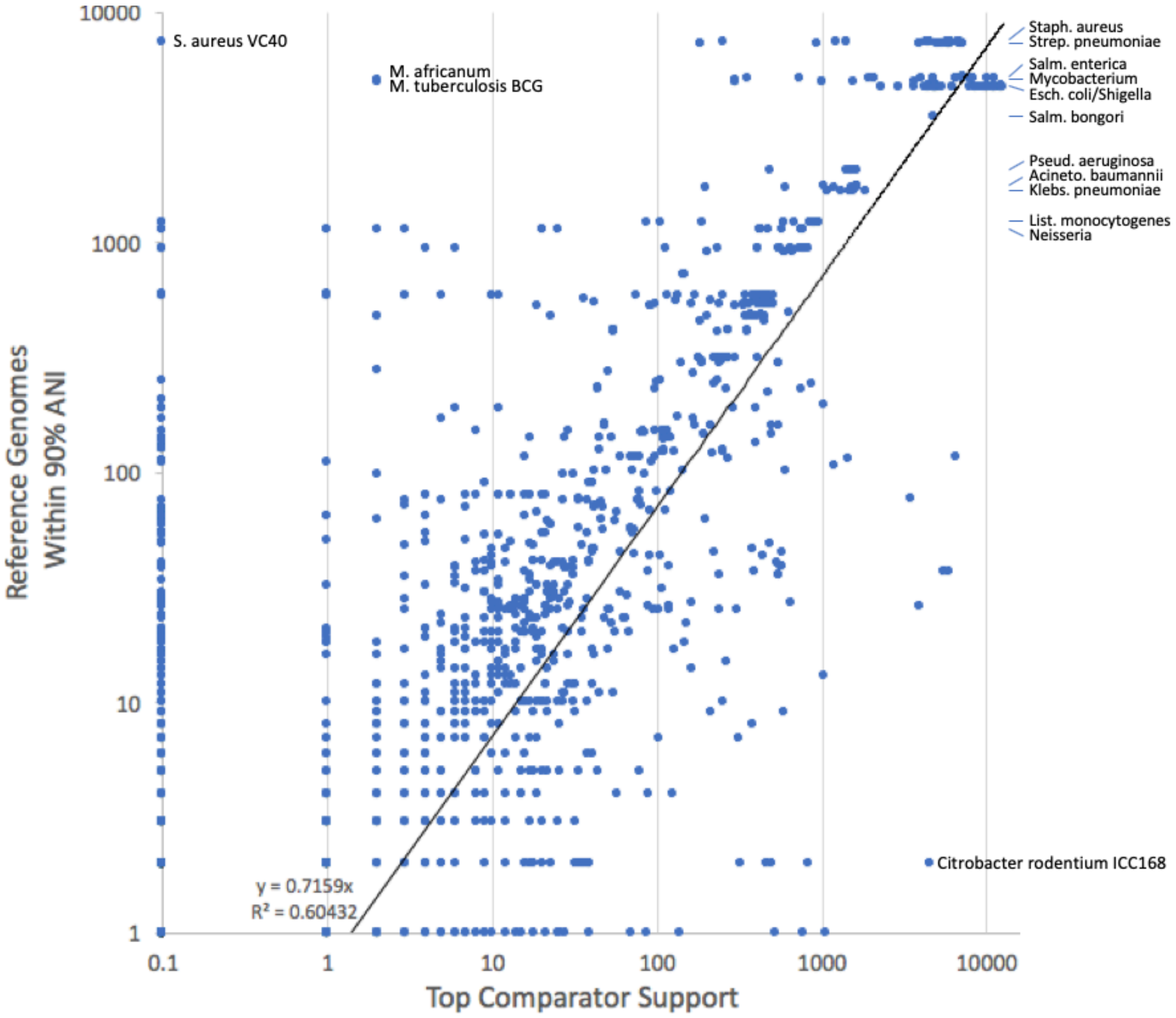

Supp. Fig. 10. Bacterial IGE typing. See Methods. The 6278 TIGER/Islander bacterial IGEs were analyzed, omitting archaeal IGEs and the corresponding mock IGEs.

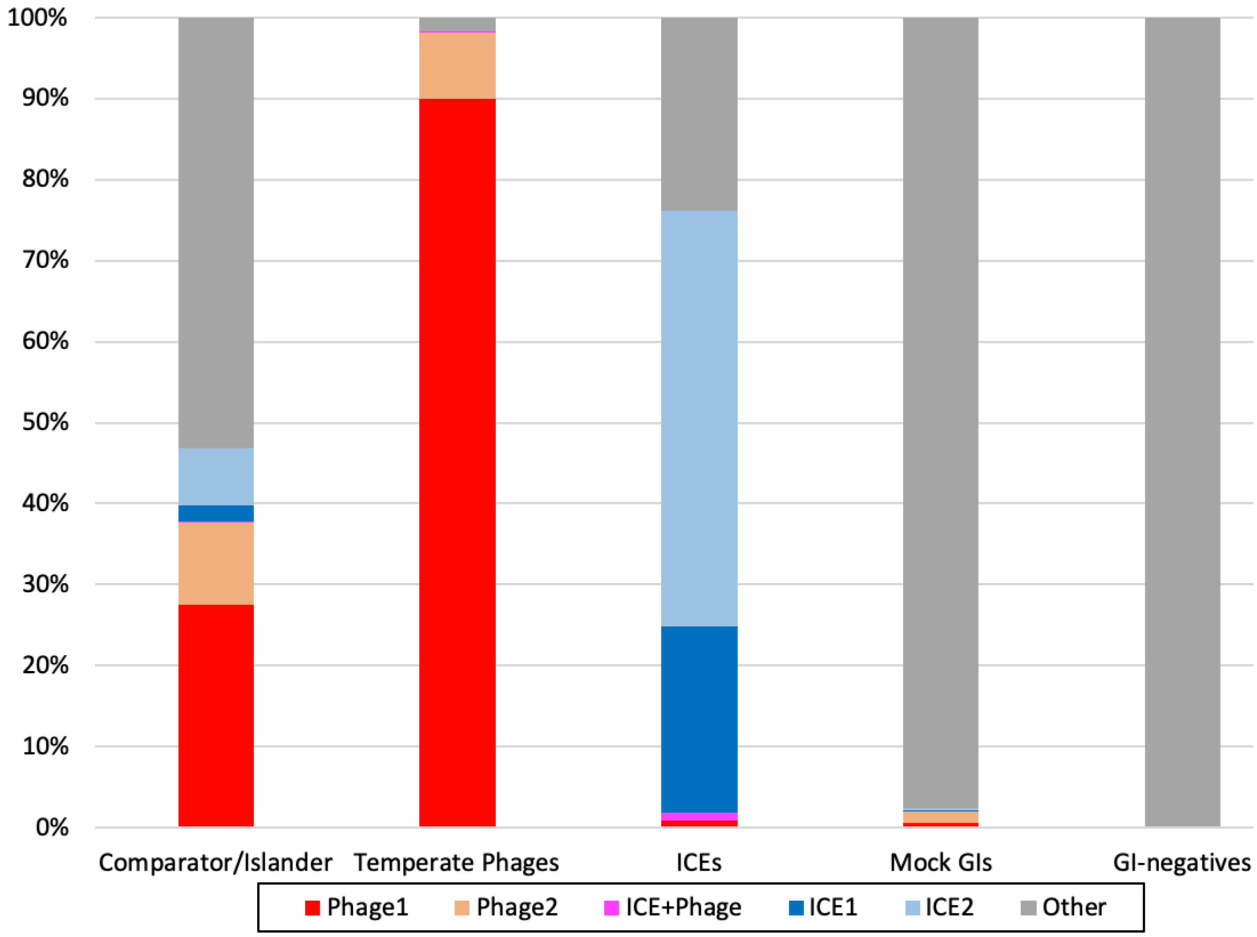

Supp. Fig. 11. Promiscuity in the Y-Int tree. Red: high support clades in the promiscuity triangles of Fig. 4; gray: low support clades; pink asterisks: Pfam domain-breaking integrases. Scale bars shown for the full tree and all insets.

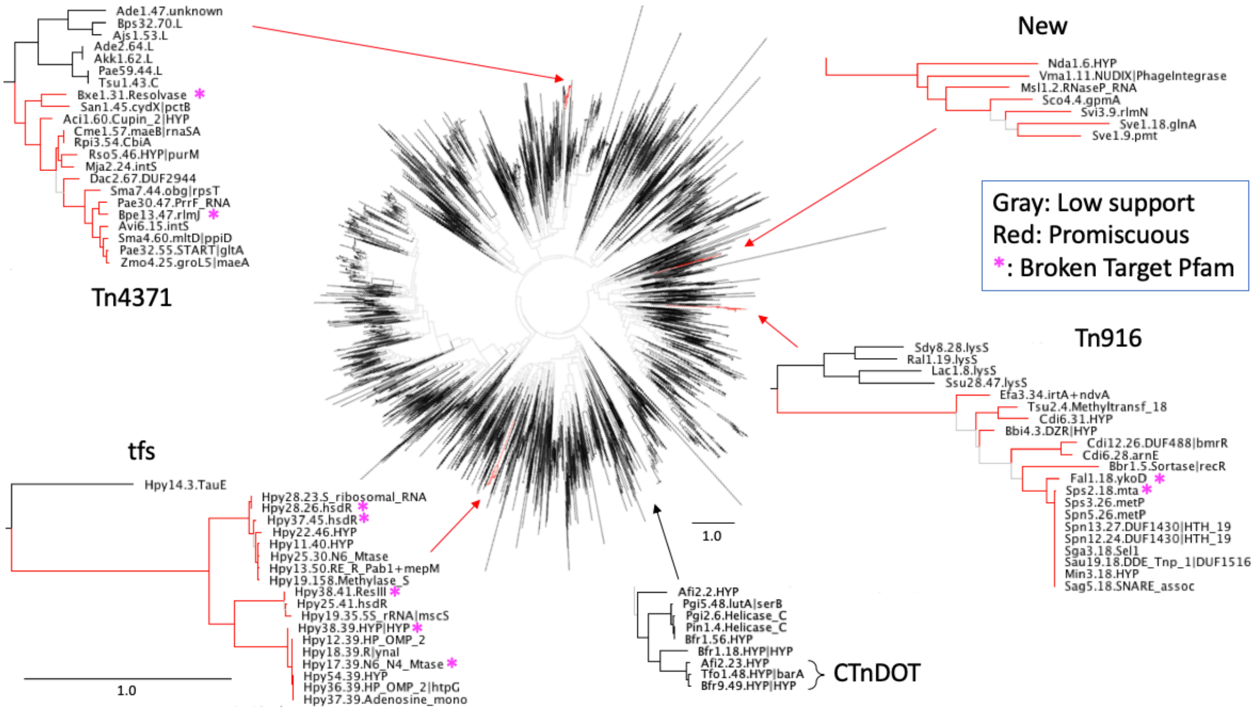

Supp. Fig. 12. Promiscuity in the S-Int/S-Core tree. Red: high support clades in the promiscuity triangles of Fig. 4; copper: high support S-Int clades; blue: high support S-Core clades; gray: low support clades; pink asterisks: Pfam domain-breaking integrases. Scale bars shown for the full tree and all insets.

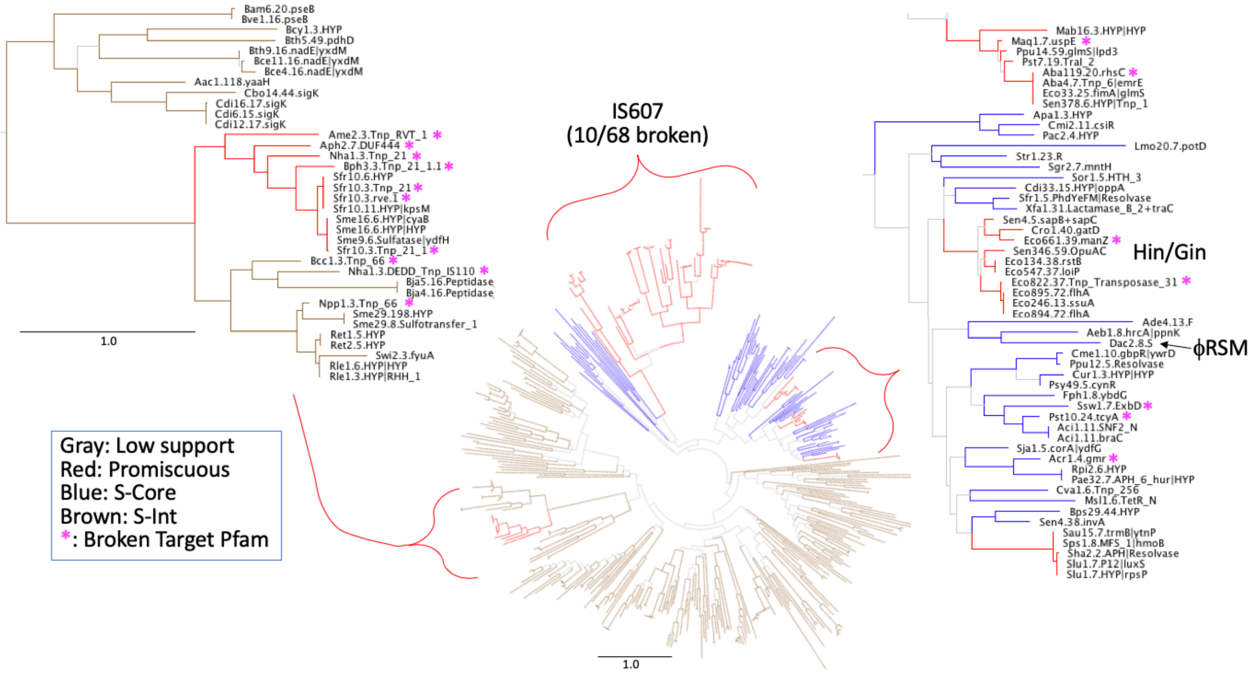

SUPPLEMENTARY FILES

Supplementary File 1. Islander/TIGER IGEs from 2168 prokaryotic genomes.

Supplementary File 2. Phylogenetic tree of IGE-unique Y-Ints. In FigTree format; see Supp. Fig. 8 for color coding.

Supplementary File 3. Phylogenetic tree of IGE-unique S-Ints and S-Cores. In FigTree format; see Supp. Fig. 8 for color coding.

Supplementary File 4. IGE-broken genes. The 410 TIGER IGEs that broke Pfam domains (see Table 1).
