## Supplementary material for "New candidates for regulated gene integrity revealed through precise mapping of integrative genetic elements": Supp. Table 11

Sau359.24.rlmH supp=173 type=Other int=S-Int,S-Int islander=no lit=SCC(23911) (mapped by Ito et al)

CGGGTTGTGTTAATTGAACAAGTGTACAGAGCATTTAAGATTATGCGTGGagaagcatatcataaatgatgcggttTTTTCAGCCGCTTCATAAAGGGATTTTGAATGTATCAGAACATATGAGGT

CGGGTTGTGTTAATTGAACAAGTGTACAGAGCATTTAAGATTATGCGTGGagaggcgtatcataagtaatgaggttCATGATTTTTGACATAGTTAGCCTCCGCAGTCTTTCAAGTAAATAATATC

TATAAATTTCTGTAGTTATTTTTCAAAAACCGCATCATTAACTGATAAGCagaggcgtatcataagtaatgaggttCATGATTTTTGACATAGTTAGCCTCCGCAGTCTTTCAAGTAAATAATATC

Sau359.14.Z supp=123 type=ICE1 int=Y-Int islander=yes lit=vSa3(15)

TAAACTACACACGTAGAAAGATGTGTATCAGGACCTCTGGACGCGGGTTcaaatcccgccgtctccattataaagcctgcaacccatgtggttgtaggctttttGTTTTTGGGTGCACAAAAAGTGCACAGATAGTGCACAGTACATAAAAAAG

TAAACTACACACGTAGAAAGATGTGTATCAGGACCTCTGGACGCGGGTTCaaatcccgccgtctccatatttgtagcctacaacctttgtggatgtgggctttttTATATGTTTTTTATCTATTCTTGTGAGGAAGGTTAAATTAGCTGTGTAC

TATGACAATGCTTATAGAGAGTTATACATGAATAGATAAACGCTTTAATGaactcccgccgtctccatatttgtagcctacaacctttgtggatgtgggctttttTATATGTTTTTTATCTATTCTTGTGAGGAAGGTTAAATTAGCTGTGTAC

Sau359.46.HYP supp=1419 type=Phage1 int=Y-Int islander=no lit=phiSa2(109)

GATGATATTGTTAAGAACATGAAGCCTTTGATTGTACAAATGATATTTGAaccatcacattatgatgatatgtttatttcaaGAAAAGCTTTAACGCCAGTGTTCTCAAGCGTTTTATAAAGCTTGTAAAAA

GATGATATTGTTAAGAACATGAAGCCTTTGATTGTACAAATGATATTTGAaccatcacattatgacgatatgtttattttaaACACACAAGCTCATGCGCGTCTTGATCAAATGGCACAACAGTTTGAAGTT

GGTATAACAAGGGTTTTATACATTTGCGTACAACGACGAAATGTCAATTTaccatcacattatgacgatatgtttattttaaACACACAAGCTCATGCGCGTCTTGATCAAATGGCACAACAGTTTGAAGTT

Sau359.43.hlb supp=1212 type=Phage1 int=Y-Int islander=no lit=phiSa3(70)

TACTGATTTGAAGTTAGTTAGTCATAACGTTTATATGTTATCGACCGTTTtgtatccaaactggAGACTTTTAACATAAAATTACTTATCATTCAAAAAGTAAAACAGCATAAT

TACTGATTTGAAGTTAGTTAGTCATAACGTTTATATGTTATCGACCGTTTtgtatccaaactggGGGCAATATAAACGCGCTGATTTAATCGGACAATCTTCTTATATTAAAAA

ACCCATTAGGGACTCCAAACCCAATAAATACTGTTGTTACAAGGTTTCTAtgtatccaaactggGGGCAATATAAACGCGCTGATTTAATCGGACAATCTTCTTATATTAAAAA

Spy14.40.csn2|lepA supp=45 type=Phage1 int=Y-Int,Y-Int islander=no lit=48_1(4538)

GACAAATAGTGCGATTACGAAATTTTTTAGACAAAAATAGTCTACGAGGTtttagagctatgAAAATCTTTTATTTATGATATAATGATTTGTGTGTAAATATGCGCTTCT

GACAAATAGTGCGATTACGAAATTTTTTAGACAAAAATAGTCTACGAGGTtttagagctatgCTGTTTTGAATGGTCTCCATTCAACATTGCCAATGGTAACTTGAGAAAG

GTATAGTCGGGGACAGGTAGTCCAATCTATTGATATAAAGCCTTTTTAGtttacgagctatGCTGTTTTGAATGGTCTCCATTCAACATTGCCAATGGTAACTTGAGAAAG

Spy14.41.Z supp=180 type=Phage1 int=Y-Int islander=yes lit=48_2(4786)

AACCTATGGACGTAGACAAATATGTTGGCAGGTGTTTGGACGTGGGTTCGactcccaccggctccatcaatgcttaccgtaagtaatcataacttactaaaaccttgttacatcaaggttttttctttttgccttgttcatgagttACCATAACTTTCTATATTATTGACAACTAAATTGACAACTCTTCAATTAT

AACCTATGGACGTAGACAAATATGTTGGCAGGTGTTTGGACGTGGGTTCGactcccaccagctccatcaatgcttaccgtaagtaatcataacttactaaaaccttgttacatcaaggttttttctttttgtcttgttcatgagttTCCGTTATCCCTATAGCCATACCATCACGCTTACCGATACTCCATCAATA

TAGATTGACAACTAATTCTCAACAAACGTTAATTTAACAACATTCAAGTAactcccaccggctccatcaatgcttaccgtaagtaatcataacttactaaaaccttgttacatcaaggttttttctttttgtcttgttcatgagttACCGTTATCCCTATAGCCATACCATCACGCTGACCGATACTCCATCAATA

Spy14.34.hup supp=123 type=Phage1 int=Y-Int islander=no lit=48_3(4425)

TTGAAATTGCAGCTTCAAAAGTTCCAGCCTTCAAAGCTGGTAAAGCTCTTaaagacgctgttaaataattGCATATATAAAAGCCCATTGTACAAGCGTTGTAGCCTGTGCAGTGGGCTT

TTGAAATTGCAGCTTCAAAAGTTCCAGCCTTCAAAGCTGGTAAAGCTCTTaaagacgctgttaaataattCGTCTAGAAAAATCTTGTTGCTATCGATGTTTATTGATAGCGACAAGGTT

TAAAATCAAAGTTTATTATATTATATGTGCACAATATTAAGTAATCTTTAaaagacgctgttaaataattCGTCTAGAAAAATCTTGTTGCTATCGATGTTTATTGATAGCGACAAGGTT

Spy14.42.Response_reg|msrAB supp=244 type=Phage1 int=Y-Int islander=no lit=48_4(1587)

AGGGCTATGGTTATTTATTATCATCTGTAGAATAGGGTTAGGCCTAAAAAtctgatataatataagaTTACCTCTCAGAATCATTGATATCAAGGCTTTTTTAAAGTCTAAAAACAG

AGGGCTATGGTTATTTATTATCATCTGTAGAATAGGGTTAGGCCTAAAAAtctgatataataaaagaAAGACTAGTATGAGAGGGCCATGCAAGGCCCTTTATAAAGATAAGGAGAC

TATCAAGTGATTAGGCAATGATAGCAAATGATACCATATTAATGTTCTTTtctgatataataaaagaAAGACTAGTATGAGAGGGCCATGCAAGGCCCTTTATAAAGATAAGGAGAC

Spy14.38.recX supp=243 type=Phage1 int=Y-Int islander=no lit=48_5(2941)

CAACACATTTTTAGCAGTATTTTCGTCTTCTATTTTACCATTTTTGATATaatcaaagtatagataaaactcctaaaattgTGGCTTTACAATGTCGTTCCTCGACTATTCCCCGACTTCTGGACGAG

CAACACATTTTTAGCAGTATTTTCGTCTTCTATTTTACCATTTTTGAtataatgaacatatgaaaatcactaaaattgAAAAGAAAAAACGCCTCTACCTTATCGAATTGGATAATGACGATTCCCTT

ATATCAATGTTTTAAAACAATTTGCTTGTATATAAATTTTAAAATACtataatcaaagtatgaaaatcactaaaattgAAAAGAAAAAACGCCTCTACCTTATCGAATTGGATAATGACGATTCCCTT

Spy14.40.HYP|proB supp=257 type=Phage1 int=Y-Int islander=no lit=48_6(4155)

TTTAACGAAGAGAGGTCATTATTCGGAAAAACAGTCCTCTGGGGCTGTTTtttatgctataatATCATCACACCTTAGAAATCCCTTGAAATCAATGGGTTTCTAAATCCAAA

TTTAACGAAGAGAGGTCATTATTCGGAAAAACAGTCCTCTGGGGCTGTTTtttatgctataatTTGGGTATGATGAAACGACAATTTGAAGATGTAACACGTATCGTAATTAA

TTTATAAAGTGATTGAGTAATGATAGCTATTGAGTTTAAGAGGTCTGGGAtttatgctataatTTGGGTATGATGAAACGACAATTTGAAGATGTAACACGTATCGTAATTAA

Eco160.2.HYP|HYP supp=6 type=Other int=Y-Int islander=no lit=c0253-c0368(97497)

ATTTGTATCAGCTAATGAGTACAGGTCTGAATCTAATCTGACAGTCCGCTctgtgccaggagcggacTGTCCAACCCATCCCTGAAGAGCCTAGATTATGTATGGTAATTCAGTCAT

ATTTGTATCAGCTAATGAGTACAGGTCTGAATCTAATCTGACAGTCCGCTctgtgccagatgcggacGTTGCTCTATTCATGTAAGATTAATAGAATGTCTTTCGATAAATTTTTGA

GATCTCTGAATCGATTGAAGTCTAAGGTTGCCAGGGCTGCAACAGCAGTCctgtgccagatgcggacGTTGCTCTATTCATGTAAGATTAATAGAATGTCTTTCGATAAATTTTTGA

Eco160.6.HYP|betA supp=2285 type=Other int=Y-Int islander=no lit=none(below lit’s 30 kbp cutoff)

GAATGTTCAGGGTAAATATCAGCAAAAAGCCAGCATCATGAATACTGGATatTTATAGCCTGCAGGCAGCATACTGCTGCCTGCAGGGGGTATAACAGTCGA

GAGTGTTCGGGTTAAATATCAGCAAAAAACCAGCATCATGAATACTGGATatGAAGCATGAGAGTTACCTCAGTGTTTTATATAAGGATTCGGTCCTCATAG

TCAGCCAGATTGAGCAAGTCAGCTTAGAATTTGCTGGCGAGATTGTCACGatGAAGCATGAGAGTTACCTCAGTGTTTTATATAAGGATTCGGTCCTCATAG

Eco160.33.RybB_RNA supp=4479 type=Phage1 int=Y-Int islander=no lit=intT-ogrK(540)

TGTCCCCATTTTGTGGAGCCCATCAACCCCGCCATTTCGGTTCAAGGTTGatgggttttttgttgcctgaaatttatGCCGTTTAAAATCATGATGTTAGAAGCACTGTTTTTTAACGATGGCGACA

TGTCCCCATTTTGTGGAGCCCATCAACCCCGCCATTTCGGTTCAAGGTTGatgggttttttgttgcctgaaatttatCTACTACCCAGCAATCCTCTCAACCATCCTCAAAATCTCCTCGCGTGATA

ACTTATTGTTTTTACTCTAACTTATTGTTTTCATTAACCCGTTTACATAAatgggttttttgttgcctgaaatttatCTACTACCCAGCAATCCTCTCAACCATCCTCAAAATCTCCTCGCGTGATA

Eco160.114.S supp=677 type=Other int=Y-Int islander=yes lit=c1165-c1293(483)

GAGCACGCCTGGAAAGTGTGTATACGGCAACGTATCGGGGGTTCGAATCCccccctcaccgccagattACAAGCGCATGTTATTGTTTGACATGCGCTTTTTTATTTATCCTTCCAGA

GAGCACGCCTGGAAAGTGTGTATACGGCAACGTATCGGGGGTTCGAATCCccccctcaccgccatattTAAAGAAGAGCTCGTACGAAAGTACGAGCTTTTTTTTCGTATATTGCATA

AATGAAATTATATAACCCCCCTGGATTAAGTCAGATTTATTTCAGGCGGTccccctcaccgccatattTAAAGAAGAGCTCGTACGAAAGTACGAGCTTTTTTTTCGTATATTGCATA

Eco160.45.potB supp=4859 type=Phage1 int=Y-Int islander=no lit=c1400-c1474(735)

TATCGTAATGGGCCTGATGTTGCTGGTTTACTGGCGCGCTTCTCGTCTGCtgaataagaaggtgAGCGAATTAGATGATTAATCCTTTATATTCAATGCATTAAGTTTGTTTTT

TATCGTAATGGGCCTGATGTTGCTGGTTTACTGGCGCGCTTCTCGTCTGCtgaataagaaggtgGAACTCGAATGATCGGTCGACTGCTTCGCGGCGGTTTTATGACCGCTATC

AGAAAGGGAAAAATTGAATAAATTCAAAATCCTGAAAGTGTCCCGCCTTAtgaataagaaggtgGAACTCGAATGATCGGTCGACTGCTTCGCGGCGGTTTTATGACCGCTATC

Eco160.11.phoQ supp=4264 type=Phage1 int=Y-Int islander=no lit=none(below lit’s 30 kbp cutoff)

GCGGATGGAAGTGATTTTTGGTCGCCAGCATTCTGCACCGAAAGATGAATaaatatgtccTTTTATCACTACATCAAGGCAAGCCATTGATTTAATTAAATGATTTGCAC

GCGGATGGAAGTGATTTTTGGTCGCCAGCATTCTGCACCGAAAGATGAATaaatatgtccATACTTCACGCATTACGTTAAGCATCCGTTATAATCGGTTGCAGATACCA

ATAGTGAACAATCTGAAACATTATGAAACGGCAAAAAACACTAATTGATAaaatatgtccATACTTCACGCATTACGTTAAGCATCCGTTATAATCGGTTGCAGATACCA

Eco160.55.icd supp=149 type=Phage1 int=Y-Int islander=no lit=c1518-c1601(669) (crossover region from Wang 1997)

CCCAGGCTCTATTATTCTCTCCGCTGAGATGAtgctgcgccatatgggctggactgaagccgcagacctgattgttaaaggtatggaaggcgcaatcaatgcgaagaccgtaacctatgacttcgaacgtctgatggaaggcgctaagctgctgaaatgttcagagtttggtgatgcgatcatcaagaatatgtaatcactacatgtgttaaatattgtaacgggcgtataacacgcccgttgttttATGATGATGTAAAATCTTCCCCAAAACCCTTC

CCCAGGCTCTATTATTCTCTCCGCTGAGATGAtgctgcgccacatggggtgggttgaagcggctgacttaattgttaaaggtatggaaggcgcaatcaacgcgaagaccgtaacctatgacttcgaacgtctgatggaaggcgctaaactgctgaaatgttcagagtttggtgacgcgatcatcaagaacatgtaatctctacatgtgttaaatattgaaacgggcgtataacacgcccgttgttttTATAAATATGTTAACCGTTATAAAATAACGTA

ATCAGCGAGTTGAATACATAATTTTTATATACtgctgcgccacatggggtgggttgaagcggctgacttaattgttaaaggtatggaaggcgcaatcaacgcgaagaccgtaacctatgacttcgaacgtctgatggaaggcgctaaactgctgaaatgttcagagtttggtgacgcgatcatcaagaacatgtaatctctacatgtgttaaatattgaaacgggcgtataacacgcccgttgttttTATAAATATGTTAACCGTTATAAAATAACGTA

Eco160.23.S supp=1173 type=Other int=Y-Int islander=yes lit=none(below lit’s 30 kbp cutoff)

ACCGGTCTCGAAAACCGGAGTAGGGGCAACTCTACCGGGGGTTCAAATCCccctctctccgccaATCATTCAACAAAATCAATCACTTACAAAGCATTTTTGATTTTGTCTGCA

ACCGGTCTCGAAAACCGGAGTAGGGGCAACTCTACCGGGGGTTCAAATCCccctctctccgccaCTTTATCAATGACTTATCTCCCGACTTCCCGCCTTGCTTTTCCTAAACAG

AAAGGAAAAATAGCACTTTGGTGGGCATTGTCGAACACTAAAAATTTAGAccctctctccgccaCTTTATCAATGACTTATCTCCCGACTTCCCGCCTTGCTTTTCCTAAACAG

Eco160.31.DUF2545 supp=1 type=Other int=Y-Int,Y-Int islander=no. XXX

CCTCCTAAGGGCTAATTGCAGGTTCGATTCCTGCAGGGGACACCATCTATCATCCCCCTCGCAATAACCCACTTACAATCAGTGAGTTATCTTAAAATCC

CCTCCTAAGGGCTAATTGCAGGTTCGATTCCTGCAGGGGACACCATTTATTACCGCCTTCCATACCCCACCATAAAGCGGATCATCAAGCCACAAGCTGG

ATGTCTGATGCTGATGTGAACACCTGCCCAAATGAGCCATCCTGACCCACTATCGCCTGCCATACCCCACCATAAAGCGAATCATCAAGCCACAAGCCGG

Eco160.48.Z supp=752 type=Phage1 int=Y-Int,S-Core islander=yes lit=c3143-c3206(625)

AAGACTYACTAAGCATGTAGTACCGAGGATGTAGAAATTTCGgacgcgggttcaactcccgccagctccaccaatcatyattgctcggtgtaaggacygcwccaacaaaaacaggatgttagcagtctcagcaggaccagggccaggcggtgaagggacaaaaaaggatacgcaaaggagccgcggctctttagttacatgaaagcccgctaatgcgggcttttttTATGGGATAAGACATATGAACCAACTGCAAGAAATCATTC

AAGACTGACTAAGCATGTAGTACCGAGGATGTAGGAATTTCggacgcgggttcaactcccgccagctccaccaatcatgattggacggtgtaaggactacaccaacaaaaacaggaagttataagtctcagcaggacaccgaccagacggtgaggagacaaaaaaggatacgcaacggagtcgcgactccctatgacattaaagcccgctgatgcgggcttttttATTGGTCCTGACCCGCCCTGAATAGAATTCATCCAATTACA

ACAATATTTGGCTATTTAAATTCAATGAATTACATAACTATggacacgggttcaactcccgccagctccaccaatcatgattggacggtgtaaggactacaccaacaaaaacaggaagttataagtctcagcaggacaccgaccagacggtgaggagacaaaaaaggatacgcaacggagtcgcgactccctatgacattaaagcccgctgatgcgggcttttttATTGGTCCTGACCCGCCCTGAATAGAATTCATCCAATTACA

Eco160.105.F supp=166 type=Other int=Y-Int islander=yes lit=c3556-kpsM(18611)

GGATAGCTCAGTCGGTAGAGCAGGGGATTGAAAATCCCCGTGTCCTTGGttcgattccgagtccgggcaccaAATTCATATAAACGGACCTCCACGGAGGTCCGTTTTTCGTTTCAGAACGC

GGATAGCTCAGTCGGTAGAGCAGGGGATTGAAAATCCCCGTGTCCTTGGttcgattccgagtccgggcaccaCTA-TTCTAAAGAACCCGCCCACAAGGCGGGTTTTTGCTTTTGGATCTTT

AACAACAAATACAAAATAATCTATATAAAACAATTTGTTATACAATCAGTttcattccgattccgggcaccacTAATTCTTAAGAACCCGCCCACAAGGCGGGTTTTTGCTTTTGGATCTTT

Eco160.52.F supp=2986 type=Other int=Y-Int islander=yes lit=c5143-c5216(318)

CGGATAGCTCAGTCGGTAGAGCAGGGGATTGAAAATCCCCGTGTCCTTGGttcgattccgagtccgggcaccaaattcatatcaacggacctccacggaggtccgtttttcgtttcagaacgccacgatttaagcgttctgcctacaaatcaattctaccgaactcaaccagattccccccacatcaACCCCATTGTGTGGGTATAATTGCGGGCATACCCCAGTTCGACAGAATTT

CGGATAGCTCAGTCGGTAGAGCAGGGGATTGAAAATCCCCGTGTCCTTGGttcgattccgagtccgggcaccaaattcatataaacggacctccacggaggtccgtttttcgtttcaggacgccacgatttaagcgtcctgccgccaaatcaattctaccgaactcaaccagattctccccacatcaCCAGCAATTTGCGGGCATATCCCAATTCGGGAAAATTTGTTTCTGAGCTA

TTTAATAAAACCAAATAAATTATATTAATATCAATATGATAAAAGAAATTttcgattccgagtccgg-caccaaattcatataaacggacctccacggaggtccgtttttcgtttcaggacgccacgatttaagcgtcctgccgccaaatcaattctaccgaactcaaccagattctccccacatcacCAGCAATTTGCGGGCATATCCCAATTCGGGAAAATTTGTTTCTGAGCTA

Eco661.11.T supp=869 type=Phage1 int=Y-Int,S-Core islander=yes lit=87_1(50)

GTGAACATGTCCTT-TCAGGGCCGATATAGCTCAGTTGGTAGAGCAGCGCattcgtaatgcgaaggtcgtaggttcgactcctattatcggcaccatttAAATCAATAAGTTACACATCATTAGTACCTTCCTTATTTTTTGACTGGGA

GTGAACACACCCTTCTCAGGGCCGATATAGCTCAGTTGGTAGAGCAGCGCattcgtaatgcgaaggtcgtaggttcgactcctattatcggcaccatttCAACGTCCCAAAACGTCCGTATTCATCCATAAATACCCTGATTTATAAAG

GAGCGCAGTGTTGATGGGGTAATGCTTTGAATTAGAAGCGGATTCTTATAattcgtaatgcgaaggtcgtaggttcgactcctattatcggcaccatctCAACTTCCTCAAACGTCCGTATTAGTCCATAAAATCTCTGATTTATAACA

Eco661.13.T supp=1 type=Phage2 int=Y-Int islander=no lit=87_2(44)

AATTAGAAGCGGATTCTTATAATTCGTAATGCGAAGGTCGTAGGTTCGACtcctattatcggcaccatCTCAACTTCCTCAAACGTCCGTATTAGTCCATAAAATCTCTGATTTATAA

AATTAGAAGCGGATTCTTATAATTCGTAATGCGAAGGTCGTAGGTTCGACtcctattatcggcaccatTTAAATCAAGTAGTTACCCCATATTCAAATACACCACGTTTCCTCCTGTG

TCTATTTTAATTTATGTATTAAAATCATTGTGTTGGTGGCGATTATTGGTtcctattatcg-caccattTAAATCAAGTAGTTACCCCATATTTAAATACACCACGTTTCCTCCTGTG

Eco661.39.PBP|ybhC supp=2468 type=Phage1 int=Y-Int islander=no lit=87_3(11)

CTTAAAGGTATTAAAAACAACTTTTTGTCTTTTTACCTTCCCGTTTCGCTcaagttagtataaaaaagcTGAACGCGAAACAGCAAAACCCAATAATATCAATGTATTACAATGCATTT

CTTAAAGGTATTAAAAACAACTTTTTGTCTTTTTACCTTCCCGTTTCGCTcaagttagtataaaaaagcAGGCTTCAACGGATTCATTTTTCTATTTCATAGCCCGGAGCAACCTGTGA

AGTGACCTGTTCGTTGCAACAAATTGATAAGCAATGCTTTTTTATAATGCcaacttagtataaaaaagcAGACTTCAACGGATTCATTTTTCTATTTCATAGCCCGGAGCAACCTGTGA

Eco661.50.S supp=3485 type=Phage1 int=Y-Int islander=yes lit=87_4(116)

TGTTTATAGTGCGCGTCATTCCGGAAGTGTGGCCGAGCGGTTGAAGGcaccggtcttgaaaaccggcgacccgaaagggttccagagttcgaatctctgcgcttccgccaaataa-tcaaggggtta-ccaaatggtagccccttt-gcttttTTGGCTCTTGGTATAAACTTGGTATACCAATTGGTATATTTTTATTCGGA

TGTTTATAGTGCGCGTCATTCCGGAAGTGTGGCCGAGCGGTTGAAGGCACcggtcttgaaaaccggcgacccgaaagggttccagagttcgaatctctgcgcttccgccagataagataaggggttagctaaatgctaacccctttttcttttGCCTGTCGAAATTCTCAGGGCGTTATATTTGCTTAATGACCTGATAATCC

TAACTTTTCAGCAATTTCACGTATTACACTGATTTTTATTTGTTTTTCTTcggtcttgaaaaccggcgacccgaaagggttccagagttcgaatctctgcgcttccgccagataagataaggggttagctaaatgctaacccctttttcttttGCCTGCCGAAATTCTCAGGGCGTTATATTTGCTTAATGACCTGATAATCC

Eco661.63.wrbA supp=4325 type=Phage1 int=Y-Int islander=no lit=87_5(9)

GGAGTGGTTAGAAATGGCTAAAGTTCTGGTGCTTTATTATTCCATGTACGgacatattgaaaccatgAAGCATTAAAAACACAGAATTACCGAAGATCGTCCAAGAGCATATATTTT

GGAGTGGTTAGAAATGGCTAAAGTTCTGGTGCTTTATTATTCCATGTACGgacatattgaaacgatgGCACGCGCAGTCGCTGAGGGTGCAAGCAAAGTGGATGGCGCTGAAGTTGT

AGGAGGATGAAATAAAATTATATTTTTCAAAATGATAAACCATATCCTTAgacatgttgaaacgatgGCACGCGCAGTCGCTGAGGGTGCAAGCAAAGTGGATGGCGCTGAAGTTGT

Eco661.86.S supp=326 type=Other int=Y-Int islander=yes lit=87_6(22)

AGCACGCCTGGAAAGTGTGTATACGGCAACGTATCGGGGGTTCGAATCCCcccctcaccgccagattataaGCGCATGTTATTGTTTAACATGCGCTTTTTTATTTTCCCTTCCAGACAG

AGCACGCCTGGAAAGTGTGTATACGGCAACGTATCGGGGGTTCGAATCCccccctcaccgccatatttaaAGAAGAGCTCGTACGAAAGTACGGGCTTTTTTTTCGTATATTGCACACAC

ATGAAATTATATAACACCCCTGGATTAAGTCAGATTTATTTCAGGCGGTccccctcaccgccatatttaaAGAAGAGCTCGTACGAAAGTACGGGCTTTTTTTTCGTATATTGCACACAC

Eco661.48.potB supp=4726 type=Phage1 int=Y-Int islander=no lit=87_7(12)

TATCGTAATGGGCCTGATGTTGCTGGTTTACTGGCGCGCTTCTCGTCTGCtgaataagaaggCAACAGAAATAGATAATTAGTGCTTTATATTCAAAAGGTTAATAGGAAAT

TATCGTAATGGGCCTGATGTTGCTGGTTTACTGGCGCGCTTCTCGTCTGCtgaataaaaaggTGGAACTCGAATGATCGGTCGACTGCTTCGCGGCGGTTTTATGACCGCTA

AGAAAGGAAAAAATTGAATAAATTCAAAATCCTGAAAGTGTCCCGCCTTAtgaataaaaaggTGGAACTCGAATGATCGGTCGACTGCTTCGCGGCGGTTTTATGACCGCTA

Eco661.15.phoQ supp=4224 type=Phage1 int=Y-Int islander=no lit=87_8(10)

GCGGATGGAGGTGATTTTTGGTCGCCAGCATTCTACGCCGAAAGATGAATaaatatgtccTTTTATCACCACAGCAAGGCAACCCATTGATTTAATTAAACAACCCACAC

GCGGATGGAGGTGATTTTTGGTCGCCAGCATTCTACGCCGAAAGATGAATaaatatgtccATACTTCACGCATTACGTTAAGCATCCGTTATAATCGGTTGCAGATACCA

ATAATGAACAATCTGAAACATTATGAAACGGCAAAAAACACTAATTGATAaaatatgtccATACTTCACGCATTACGTTAAGCATCCGTTATAATCGGTTGCAGATACCA

Eco661.58.ompW supp=2204 type=Phage1 int=Y-Int,Y-Int islander=no lit=87_10(121)

GCAGGTGGGGGTTGATTATCTGATTAACCGTGACTGGTTGGTTAACATGTcagtgtggtacatggatatcgataccacggctaagtacaaatctggtgttactaccgtaaaagactcggtacgactggatccgtgggtgtttatgttctccgcaggatatcgtttttaattTTCTCTGCAAAACCTCTGCAAAACCCCTCTGCAAAACTGGTCATCAAATG

GCAGGTGGGGGTTGATTATCTGATTAACCGTGACTGGTTGGTTAACATGTcagtgtggtacatggatatcgataccaccgccaattataaactgggcggtgcacagcaacatgatagcgtacgcctcgatccgtgggtgtttatgttctcagcaggatatcgtttttaattCCGCACAAAAACGACCCCGTAATATACGGGGTCAATAAGGACATGGCATA

CTGTATTTTTCCTTTAAATATAATATGTTATTGAGTTTTGTAAAATCTACcagtgtggtacatggatatcgataccaccgccaattataaactgggcggtgcacagcaacatgatagcgtacgcctcgatccgtgggtgtttatgttctcagcaggatatcgtttttaattCCGCACAAAAACGACCCCGTAATATACGGGGTCAATAAGGACATGGCATA

Eco661.51.ttcA supp=2219 type=Phage1 int=Y-Int islander=no lit=87_11(43)

GCATACTCTTCTGATGCCATATAACGAATTGAGTCGCTTTTAAATGTCGCaaaatcaagaaattagcaagaaagaacaatacaacctgaacaaGTCAGAGTAAAAACCGCAACTATCTGACGAACAATAACTTTTATGTTTTT

GCATACTCTTCTGATGCCATATAACGAATTGAGTCGCTTTTAAATGTCGCaaaatcaagaaattagcaagaaagaacaatacaacctgaacaaATTACAAAAACGCCTGCGTCGTAACGTGGGCGAAGCCATTGCTGACTTCA

TTTGCCCCCAATTTGCCCCATTTAGTACCAGAGAACTGAAATAATGCAAGaaaatcaacaaattacaaagaaagaacaatacaacctgaacaaATTACAAAAACGCCTGCGTCGTAACGTGGGCGAAGCCATTGCTGACTTCA

Eco661.21.L supp=3692 type=Phage1 int=Y-Int islander=yes lit=87_14(236)

CCTCCACTTTCTTCCCGAGCCCGGATGGTGGAATCGGTAGACACAAGGGAtttaaaatccctcggcgttcgcgctgtgcgggttcaagtcccgctccgggtaccatgggaaagctaagaataaaatcaaagcaataagcagtgtcgtgaaaccaccttcgggtggttttttcgtATTTGTATTTTGTGCAATGGCGGTAGAGTGGCGATGGTGTGGCGATATGC

CCTCCACTTTCTTCCCGAGCCCGGATGGTGGAATCGGTAGACACAAGGGAtttaaaatccctcggcgttcgcgctgtgcgggttcaagtcccgctccgggtaccatgggaaagctaagaataaaatcaaagcaataagcagtgtcgtgaaaccaccttcgggtggtttttttgtGCCTGCAACTTGTCGTTACACCCTCCTTGATTTTTAATCACCGGCAAAGC

AAAGTTAATAAAATCATAAGGTTATGATTTATAAATGGTGGTTGATAGTTtttaaaatccctcggcgttcgcgctgtgcgggttcaagtcccgctccgggtaccatgggaaagataagaataaaatcaaagcaataagcagtgtcgtgaaaccaccttcgggtggtttttttgtGCCTGCAACTTGTCGTTACACCCTCCTTGATTTTTAATCACCGGCAAAGC

Eco661.44.S supp=2855 type=Phage1 int=Y-Int islander=yes lit=87_15(110) tRNA?

CATCATGGCAACCATCTGAACGGAGAGATGCCGGAGCGGCTGAACGGACcggtctcgaaaaccggagtgggggcaactccaccgggggttcaaatccccctctctccgccaaaat-tcaatcacttaCACATCATTAAGTCAGTGACAAAAATCACACTTGGAATTACTTGGAATAT

CATCATGGCAACCATCTGAACGGAGAGATGCCGGAGCGGCTGAACGGACCggtctcgaaaaccggagtaggggcaactctaccgggggttcaaatccccctctctccgccactttatcaatgacttaTCTCCCGACTTCCCGCCTTGCTTTTCCTAAACAGAACAATCGTAGAATAT

TTATTCCAAATAAAACACATAATATATTGATTTTATTAACCCATAGCATAggtctcgaaaaccggagtaggggcaactctaccgggggttcaaatccccctctctccgccactttatcaatgacttaTCTCCCGACTTCCCGCCTTGCTTTTCCTAAACAGAACAATCGTAGAATAT

Eco661.48.ycgE supp=3889 type=Phage1 int=Y-Int islander=no lit=87_17(25)

ATGACCGCCGTCTGTCCGTTGCGGTTTCAGCAATCCGTAACGCCTCTGCCacgcgcgtaacgtgacagggattaaGGGTAAACTAATTGATTTTGCAAGTTTTTATTTTACCCTGCTTTCTTAT

ATGACCGCCGTCTGTCCGTTGCGGTTTCAGCAATCCGTAACGCCTCTGCcacgcgcgtaacgtgacagggttaaTATCACAAAGCAACGCCACTTCACCAATTGTGTAAAGCGCCATCGTCTCA

AAAAACGGCGCTCCTGGACGATCTTCGTCGATTTTTAAAAATGTTGCGTcacgcgcgtaacgtgacagggttaaTATCACAAAGCAACGCCACTTCACCAATTGTGTAAAGCGCCATCGTCTCA

Eco661.8.R supp=5 type=Other int=Y-Int,S-Core islander=no lit=87_18(354) L end (3193327 3201523) is ~350 bp downstream of trna (3192908-3192982) within int gene XXX bad crossover

TGACGGTAAGGAGAAAATCCTGACCGTCGGAAAATATCCGCTTATGACTTtgcaagaggcaaGGGATAAAGCATGGACTGCGAGGAAAGACATCTCGGTTGGCATCGATCCG

TGACGGTAAGGAAAAAATACTGACCGTAGGAAAATATCCGTTTATGACTTtgcaggagacaaCAGATACCCTTCAAACGAAATCTACCTTCACCCCGTAAAAGATGGGTTTG

CGAAAGAGAAATAAACTACAATTCAAATTGTTAGCTAATTTTTCGATTCCtgcagg-gacaccAGATACCCTTCAAACCAAATCTACCTTCATCCCGTAAACGATGGGTTTG

Eco661.24.Z supp=644 type=Phage1 int=Y-Int,Y-Int islander=no lit=87_19(7) XXX bad crossover

TAGTACCGAGGACGTAGGAATTTCGGACGCGGGTTCAACTCCCGCCAGCTccaccaCTTTAAGAAGGACTACAACCGGACAGTAGCAATAAATACAGCCACTTAC

TAGTACCGAGGACGTAGGAATTTCGGACGCGGGTTCAACTCCCGCCAGCTccaccaATCATGATTGGACGGTGTAAGGACAACACCAACAAAAACAGGAAGTTAG

AGCAGTTATCTAGAAAATTCAATCAAATAGATATCTAGTTACAGGGACGccagccAATCATGATTGGACGGTGTAAGGACAACACCAACAAAAACAGGAAGTTAG

Eco661.8.U|espF supp=139 type=Other int=Y-Int islander=no lit=87_21(35910)

GACTCCTGTGATCTTCCGCCAGTTTGCCTCTCCTGAACTCTCCCGACATCaacaaagccccttataatcccttatatatcaggttttctttgtatcctgaggtcaaacgactttaacccatatcaagttgtttgtgggggtgcattgggggtatCAGGTTGGTTACTGGCGGAAATTCCCCCAGGAGGATGGATTCATCAAAAG

GACTCCTGTGATCTTCCGCCAAAATGCCTCTCCTGAACTCTCCCGACGTTaacaaaaccccttctaatcccttatatatcaggctttctttgtgtccagagatcaatcgactcaacctgtatcaagttgcttttggggatactttagggggtatTATTTCCTCTTGAAATATTAGCCTCCCATAATAGGAAAAAGGTAATACAT

CCAGCTGAATAACACATCCCCGTCAATACGGCCCTCGCTGTACGCTTACCaacaaaaccccttttaatcccttatatatcaggctttctttgtgtacaggggtcaaccgactcaacctgtatcaagttgcttttgaggatactttagggggtatTATTTCCTCTTGAAATATTAGCACCCCATAATGGGAAAAAGGTAATACAT

Eco661.39.manZ supp=3450 type=Phage1 int=S-Core islander=no lit=87_22(10)

TCAAGCGACTTTACCCAGTAAAAGAGGATCAGGTTGCGGCGCTCAGGCGAcacctggtgttcttTATATAGCGCGGCGATTTTTTTGATACCGATTTGCATTTCATTTGATACC

TCAAGCGACTTTACCCAGTAAAAGAGGATCAGGTTGCGGCGCTCAGGCGAcacctggtgttcttCAATACCACGCCAGCCGTATGTGGCCCGGTCATCGGCGTCACTGCCGCCA

CAGGTGATTTCACCTCCTTTCACTAAAACCCTCGTTGGTATCAAATGATTcacctggttacattTAATACCACGCCAGCCGTATGTGGCCCGGTCATCGGCGTCACTGCCGCCA

Eco661.10.L supp=500 type=Other int=Y-Int,S-Core islander=yes lit=87_23(23)

CGAAATCGGTAGACGCAGTTGATTCAAAATCAACCGTAGAAATACGTGCCggttcgagtccggccttcggcaccaaGACTCATCTAAACTATTGTTTTAAAACAATTTAATTCATAATCTTGCTTG

CGAAATCGGTAGACGCAGTTGATTCAAAATCAACCGTAGAAATACGTGCCggttcgagtccggccttcggcaccaaAAGTATGTAAATAGACCTCAACTGAGGTCTTTTTTTATGCCTGAAATCCG

TTCTTTTAAAGGCTAATAGGAACGTGATTACAGCAGGTTATGGAGACTTAggttcgagtccggccttcggcaccaaAAGTATGTAAATAGACCTCAACTGAGGTCTTTTTTTATGCCTGAAATCCG

Sen346.12.LamB_YcsF|nei supp=1214 type=Other int=Y-Int islander=no lit=93_3(4208)

ATGTACACGTTATTGCCTGAGAGCCCGACGATTGAAGCTATTATATTACGgctgCTAAAAATTATCGTGTTCTGTTCTCTTAATGTCAGGAAGAGGGATTTATT

ATATACACGTTATTGCCTGAGAGCCCGACGATTGAAGCTATTATATTACGgctgTGGATGTTCATTATGGCGTTCGATGTGCATTCCAGATAGTCAGAGGCGGG

TGCTGCGATAATGTTATTATCACTGGATACCACCGCCTGTTTGTCCATCCgctgTGGATATTCATTATGGCGTTCGATGTGCATTCCAGACAGTCAGAGGCGGG

Sen346.44.pepN|pncB supp=6628 type=Phage1 int=Y-Int islander=no lit=93_4(355)

AAATCGCGCTACGCAGAATGTTCATCTTTTCAGGCACAAACGGCCTATTTgctacatttttataatatGTACTCATAGTTTTTAAAATCGATAAAGATCGTCCAGGAGCGCTTTAAAC

AAATCGCGCTATGCAGAATCTTCATCTTTTCAGGTACAAACGCCTTTATTgctacatttttataacatACGCAGCGCAATACCATCGACCAGAAAGGTGACATATGGTGTGATCGGGG

GGTAGCAGCGCAGTAGAAATCCTTAAATTCAAGGGGTTAGCAGTCGCATCgctacatttttataacatGGGGCACGAAATGCGCTCGACCCTAAAGACAGCTTATGGTGTGATCGGGG

Sen346.59.OpuAC supp=1 type=Phage1 int=S-Core islander=no lit=93_6(16353) XXX

AGGCTTTCAGGTTCAGCCGCAAACCAATATCGCCGCGGTGATTTCACGTAatGCGATGGTGAATAAACAAATTGATATTACCTGGGAATATACCGGTACATC

AGGCTTTCAGGTTCAGCCGCAAACCAATATCGCCGCGGTGATTTCACGTAatTGGTCAACTGGAAGCGCTGATTGAAGAACTGCTCACCTATGCGCGCCTTG

AAATGAGTGAAAATCTGACTCCGCCGGAATCACAGGCGCTCAATCGCGATatTGGTCAACTGGAAGCGCTGATTGAAGAACTGCTCACCTATGCGCGCCTTG

Sen346.45.PhageIntegrase+RecT supp=1922 type=Phage1 int=Y-Int,Y-Int islander=no lit=93_9(3591)XXX

AGATGCACAGGTTACTCATGTCTATGCCGTTGCCCGCCTTAATCGATCGCtgccgactggtaagccgaactgatttcatgatcagtgccggaatcaggaaaaatagcccaaccgggaatattcacccggatgggctgacaaagaaatttgtaaaagccagaaaaatttcaggcgttaaatgtagtgataacccaccgacatttcacaagatccgtagcctggctggtcggctgtacaaaaacgaacgcggcgaggaattcgctcaaaaactactgggccacacctcagagaacaccacgaaACTCTATCTCGATGAACGCGATAATAAAGCTTACGTGATGCTCTGATTTT

AGATGCACAGGTTACTCATGTCTATGCCGTTGCCCGCCTTAATCGATCGCtgccgactggtaagccgaactgatttcatgatcagtgccggaatcaggaaaaatagcccaaccgggaatattcacccggatgggctgacaaagaaatttgtaaaagccagaaaaatttcaggcgttaaatgtagtgataacccaccgacatttcacaagatccgtagcctggctggtcggctgtacaaaaacgaacgcggcgaggaattcgctcaaaaactactgggccacacctcagagaacaccacgaaACTCTATCTCGATGAACGCGATAATAAAGCTTACGTGATGCTCTGATTTT

CACTCAGCCTTCCTGTCGCTGGTCTACGGCTTGGTACAGTAGTTGAACAGtgccgcctggtaagccggggagattatctaatcagtgccgggattagaaaaaacagccctgacggcagcattcacccggatggcctgacaaaaaaatttgtcgcagccagaaaattaacaggtatccagttcagtgaaaacccaccaacttttcacgagatcagaagtctggctggacgattgtacaaagaaacatgtggagaagaatttgctcagcgtctacttggccacacatcggagaagacaacaaaAATGTATCTTGATGAGAGAGAAAAAACGTACTTACTGCTCTGATTTTAAC

Sen346.6.G supp=2418 type=Other int=Y-Int islander=yes lit=none

AGTTCAATGGTAGAACGAGAGCTTCCCAAGCTCTATACGAGGGTTCGATTcccttcgcccgctccacgacaCTTCTTAAAGTTCAATAACTTACTGAATCCTAAGGGATATGGCTTAAAG

AGTTCAATGGTAGAACGAGAGCTTCCCAAGCTCTATACGAGGGTTCGATtcccttcgcccgctccagacaAATATCTTCTAGTATTTACTAAATCCACGAATTTACAGCAATTTTCCGTG

TGGATTCATCAGTCTTTAAGAAACAAGAAGTTAGCTAGTTGTCCTCGGCtcccttcgcccgctccagacaAATATCTTCTAGTATTTACTAAATCCACGAATTTACAGCAATTTTCCGTG

Sen346.34.comM supp=2862 type=Phage1 int=Y-Int,S-Core islander=no lit=93_14(2001)

AAAAGAGGTGCGTGACCGGGTACGTAGTGCGATTATTAATAGCGGATATGaatttccggctaaaaaaataaccattTATGTAAACGGGTTAATGAAAACAACAAGTTAGATTAAAAACAATAAGTT

AAAAGAGGCGCGTGACCGGGTACGTAGTGCGATTATTAATAGCGGATATGaatttccggctaaaaaaataactattAACCTTGCTCCAGCCGATCTACCGAAAGAGGGCGGAAGGTATGACCTGCC

CGCCATCGTTAAAAAACAGTGTTTCTAACATCATGATTTTAAACAGCTTAaatttcaggc-aacaaaaaacccattaACCTTGCTCCAGCCGATCTACCGAAAGAGGGCGGAAGGTATGACCTGCC

Sen346.13.F supp=2 type=Other int=Y-Int islander=yes lit=93_17(13023)

CTCAGTCGGTAGAGCAGGGGATTGAAAATCCCCGTGTCCTTGGTTCGATtccgagtccgggcaccaccttTCCTTTTTTATGCTTCCCTATGAATTCCAATATGTATGTAATTCAATAAG

CTCAGTCGGTAGAGCAGGGGATTGAAAATCCCCGTGTCCTTGGTTCGATTccgagtccgggcaccactcttATCAAAGTACGTTGGTTAGGGCTTGAGAACCTTAACGGACGTTACGTGG

TCAAAAACGGATTTATGACGGAATTCAATTTAATAGATAGCTGAATGCTCccgagtccgggcaccactcttATCAAAGTACGTTGGTTAGGGCTTGAGAACCTTAACGGACGTTACGTGG

Sen346.11.L supp=7 type=Phage2 int=Y-Int islander=yes lit=93_18(29808) uninterrupted from excision after damage

-AATCGGTAGACGCAGTTGATTCAAAATCAACCGTAGAAATACGTGCCGGTtcgagtccggccttcggcaccatTAGTACTTCCAAGACCATCCGAGAAAGTCCAATTATCCCTTAAAAATCAA

AAATCGGTAGACGCAGTTGATTCAAAATCAACCGTAGAAATACGTGCCGGttcgagtccggccttcg-caccatACAAAGCTTTTTTCAAGTCTACTAACGTCTACTTTAATCCACAAAAACAG

GAACATGTGTTCTTCTTTATTTCCTTTATTTTCAATTTATTATGTTGTGTtttgagtccggccttcg-caccatACAAAGCTTTTTTCAAGTCTACTAACGTCTACTTTAATCCACAAAAACAG

Vpa12.23.I supp=268 type=Other int=Y-Int islander=yes lit=VPaI1(626)

CTATAATCGTTGACCTATTAAGGCCCCTTAGCTCAGTGGTTAGAGCAGTCgactcataatcgattggtccccagttcaagtctgggaggggccaccaaattctagaaggttagtaagcgtagtgtttactaacctttttttgtacctgacgattttGGGTCAGGTAATGGGTCAGGTAAGCCCCGATCACGCACTCTTCTCTGCCT

CTATAATCGTTGACCTATTAAGGCCCCTTAGCTCAGTGGTTAGAGCAGTCgactcataatcgattggtccccagttcaagtctgggaggggccaccaaatttcgaaaggcgtagcaggttagtacctactacgccttttttgtatctgatggttttAAGTGTTGCTATGGGTACACCGTTGGCACCTGAGCTCATCTTCTTATCCC

TTTAATAGAAATTACCAAGCATTGCATACTCATAAAAATGAACCTGGTAGgactcataatcgattggtccccagttcaagtctgggaggggccaccaaatttcgaaaggcgtagcaggttagtacctactacgccttttttgtatctgatggttttAAGTGTTGCTATGGGTACACCGTTGGCACCTGAGCTCATCTTCTTATCCC

Vpa12.6.Z supp=5 type=Other int=Y-Int,S-Core islander=yes lit=VPaI2(4049)

GACTAAGCATGTAGTACCAATGATGAAAGGTTTTCGGACGCGGGTTCAACtcccgccagctccaccaaacATCATGTAAGCCTCTGATTTTTCAGGGGCTTTCTTTTTATATATGCCTAT

GACTAAGCATGTAGTACCAATGATGAAAGGTTTTCGGACGCGGGTTCAACtcccgccagctccaccaaacGTTTGGAAAAGGCCACCTTCGGGTGGCCTTTTTTGTATCAACAATCATAT

CTTCCTTGTATATCAGAGACTTACAGAGGAAAAGGCGGCGAACTCTCGAAtcccgccagctccaccaaacGTTTGGAAAAGGCCACCTTCGGGTGGCCTTTTTTGTATCAACAATCATAT

Vpa12.6.S supp=5 type=Other int=Y-Int islander=yes lit=VPaI3(25392)

GAGTTGTCCGAGTGGCTGAAGGAGCACGCCTGGAAAGTGTGTATACGGCAacgtatcgagagttcgaatctctcactcaccgccacattctAAAGGTCTGATTTATCAGGCCTTTTTTCTTATTAAAGGCTTATTGAGGTA

GAGTTGTCCGAGTGGCTGAAGGAGCACGCCTGGAAAGTGTGTATACGGCAacgtatcgagagttcgaatctctcactcaccgccacattctTGAAGAAAGACGTCCTAGGACGTCTTTTTTTATGCCTATTATACTTCTAA

TGATCTTTATACGGCACATCGCCGTTAGGTGATGGTGAAAGCGTAGCTTTacgtatcgagagttcgaatctctcactcaccgccacattctTGAAGAAAGACGTCCTAGGGCGTCTTTTTTTATGCCTATTATATTTCTAA

Vpa12.16.S supp=268(for 22.S) type= int=Y-Int islander=no lit=VPaI3(15166)

GAGCACGCCTGGAAAGTGTGTATACGGCAACGTATCGAGAGTTCGAATCTctcactcaccgccacattcTAAAGGTCTGATTTATCAGGCCTTTTTTCTTATTAAAGGCTTATTGAGGT

GAGCACGCCTGGAAAGTGTGTATACGGCAACGTATCGAGAGTTCGAATCTctcactcaccgccacattcGAAGCCCTAGCAGAAATGCTGGGGCTTTTTCGTTTTATCAATTCGTATTT

TCGACTTGCTATAAATCATTAATATTCAAATAGATACTATGTTTGGTCGActccctccctcgcacattcGAAGCCCTAGCAGAAATGCTGGGGCTTTTTCGTTTTATCAATTAGTATTT

Vpa12.17.S supp=344 type=Other int=Y-Int,Y-Int islander=yes lit=VPaI4(403)

TGGTTGAAAGCACCGGTCTTGAAAACCGGCATACGTTAATAGCGTATCTAgggttcaaatccctatctctccgccacattCAAGCCCTTGATTTATCAAGGGCTTTTCTCGTTTTAAGACGATCAAATGT

TGGTTGAAAGCACCGGTCTTGAAAACCGGCATACGTTAATAGCGTATCTAgggttcaaatccctatctctccgccacattTAGAAAAAGCCCGCTAAGAAATTAGTGGGCTTTTTCGTATATTCAGATTT

GAAAGCTAGGAAATTCAAGAGCTTGATATATATAGGTTTTCAAAGAGATGgggttcaaatccctatctctccgccacattTAGAAAAAGCCCGCTAAGAAATTAGTGGGCTTTTTCGTATATTCAGATTT

Vpa12.16.hupA supp=634 type=Other int=Y-Int,Y-Int islander=no lit=VPaI5(608)

AGATTCAAATCGCAGCAGCTAACGTACCTGCATTCGTAGCAGGTAAAGCGctgaaagaagcaTGCAATGATTAACCACATGTTTCATTAAATGTGGATAATTTCTGAAAACC

AGATTCAAATCGCAGCAGCTAACGTACCTGCATTCGTAGCAGGTAAAGCGctgaaagaatcaGTAAACTAAGTTAGAATGCACCGAGGTATCCTGCCTCGGTGCAATTTCAT

TATGGATTTAATTTCTATGTTTTGAAATGAAACAGTTTTTATTACATAGActgaaagaatcaGTAAACTAAGTTAGAATGCACCGAGGTATCCTGCCTCGGTGCAATTTCAT

Underlines mark the longest perfect matches between attL and attR. Double underlines mark the strand crossover region when experimental models can be applied. For t(m)RNA genes, this is the anticodon loop when the identity block extends that far, based on Hauser1992, Pena1996 and Smith-Mungo1994, otherwise we mark it as centered at the 3'-most nucleotide of the T-stem based on Rajeev2006. The icd strand crossover region is based on Wang1997
